# Functional Optimization of Light-Activatable Opto-GPCRs: Illuminating the Importance of the Proximal C-terminus in G-protein Specificity

**DOI:** 10.1101/2023.01.27.525823

**Authors:** Siri Leemann, Sonja Kleinlogel

## Abstract

G-protein coupled receptors (GPCRs) are the largest family of human receptors that transmit signals from natural ligands and pharmaceutical drugs into essentially every physiological process. One main characteristic of GPCRs is their ability to specifically couple with different families of G-proteins, thereby triggering specific downstream signaling pathways. While an abundance of structural information is available on GPCR interactions with G-proteins, little is known about the GPCR domains functionally mediating G-protein specificity, in particular the proximal C-terminus, the structure which cannot be predicted with high confidentiality due to its flexibility. In this study, we exploited OptoGPCR chimeras between light-gated GPCRs (opsins) and ligand-gated GCPRs to systematically investigate the involvement of the C-terminus steering G-protein specificity. We employed rhodopsin-beta2-adrenoceptor and melanopsin-mGluR6 chimeras. We discovered a dominant role of the proximal C-terminus, dictating G-protein selectivity in the melanopsin-mGluR6 chimera, whereas it is the intracellular loop 3, which steers G-protein tropism in the rhodopsin-beta2-adrenoceptor. From the functional results and structural predictions, melanopsin and mGluR6 use a different mechanism to bRhod and b2AR to couple to a selective G-protein. Collectively, this work adds knowledge to the GPCR domains mediating G-protein selectivity, ultimately paving the way to optogenetically elicited specific G-protein signaling on demand.

## 1 Introduction

Heterotrimeric, guanine-nucleotide binding G-protein coupled receptors (GPCRs) represent the largest family of membrane receptors in humans and play a principal role in physiology and pathology by transforming extracellular signals into intracellular responses. Therefore, drugs targeting GPCRs account for the largest portion of pharmaceuticals on the global market (Hauser et al., 2017).

GPCRs can be classified into five families (Class A-F) (Rosenbaum et al., 2009) and possess a highly conserved structure characterized by seven transmembrane alpha helices (TMs) with alternating extra- and intracellular loops (ILs), an extracellular N-terminus and an intracellular C-terminal tail (C-terminus) of highly variable length. GPCRs exert their downstream signaling by binding and activating heterotrimeric G-proteins. While the Galpha subunit is the main mediator of downstream signaling and triggers second messengers, the Gbeta and Ggamma subunits can influence diverse effectors, amongst them membrane ion channels. In humans, there are at least 16 known Galpha subunits, 6 Gbeta, and 13 Ggamma subunits, which can be assembled in a large variety of different combinations (Oldham & Hamm, 2008). The Galpha subunits are subdivided into four main families, each with a specific signaling pathway: the stimulatory Gs-proteins, which activate adenylyl cyclase and thereby increase the intracellular cAMP levels; the inhibitory Gi/o family, which can be further classified into Gi, Go, Gz and Gt proteins and which contrarily decrease levels of cAMP by inhibiting adenylyl cyclase; the Gq/11 family, which acts through phospholipase and includes the Gq, G11 and G15 proteins, and finally, the G12/13 group that stimulates Rho kinases (Milligan & Kostenis, 2006; Neves et al., 2002). Since these proteins trigger distinct downstream signaling pathways, selective binding of a particular GPCR to a specific G-protein is critical for accurate signal transduction (Flock et al., 2017).

While GPCRs have been extensively studied structurally, adding immensely to our undrestanding of their workings, it remains rather elusive how they selectively activate specific G-proteins. The structural components and molecular processes of GPCR-G-protein interactions were first elucidated at the structural level in bovine rhodopsin (bRhod) (Salom et al., 2006) and the beta2-adrenoceptor (b2AR) (Rasmussen, DeVree, et al., 2011; Rosenbaum et al., 2009), whereas mGluR structures were only recently resolved (Seven et al., 2021) and the crystal structure of melanopsin remains to be elucidated. From the structural studies it became obvious that the intracellular loop 3 (IL3) (Ma et al., 2020; Rasmussen, DeVree, et al., 2011), the intracellular loop 2 (IL2) (Kang et al., 2018; Qiao et al., 2020; Seven et al., 2021; Tsai et al., 2019) as well as the C-terminus (Bertheleme et al., 2013; Seven et al., 2021; Tsai et al., 2019; Tsai et al., 2018) contact the G-protein. However, contact to the G-protein does not necessarily indicate that the domain mediates G-protein selectivity, since the multiple ionic interaction networks also comprising the GPCR core structure could mediate specific conformational differences required to accommodate a particular G-protein (Venkatakrishnan et al., 2016). In group A GPCRs, the displacement of TM6 upon activation creates a cleft between TM3, TM5 and TM6 for insertion of the G-protein’s C-terminus. IL3 as a main contact point of the G-protein’s C-terminus is therefore considered to play a major role in G-protein selectivity (Rasmussen, DeVree, et al., 2011; Tsai et al., 2019; Zhou et al., 2019). This assumption was supported by functional studies with engineered chimeric Opto-GPCRs (Airan et al., 2009; Kim et al., 2005; Siuda et al., 2015; van Wyk et al., 2015). It also became obvious that the helical extensions of TM5 and TM6, protruding intracellularly and framing IL3 are dynamically involved in G-protein binding (Rose et al., 2014; Rose et al., 2015; Xu et al., 2011), rotating and extending to provide the binding pocket for the C-terminus of the G-protein (Choe et al., 2011; Lebon et al., 2011; Rosenbaum et al., 2011; Standfuss et al., 2011; Wang et al., 2013; Xu et al., 2011). However, in mGluRs – and in particular in mGluR2 (Seven et al., 2021) – cytosolic TM6 opening was not observed, but instead the G-protein’s C-terminus was stabilized by a pocket formed by IL2 and the C-terminus of mGluR2 (Seven et al., 2021). While IL3 orchestrates receptor rearrangement, it remains unclear whether it is equally implicated in the determination of G-protein specificity in mGluR2. In particular, a C-terminal region between V826-S833 was shown to affect mGluR2’s G-protein tropism (Seven et al., 2021) (Suppl. Fig.1). Opsins and other Class A GPCRs possess a conserved amphipathic helix 8 (H8) that folds into an α-helix and is arranged parallel to the cell membrane with a distal palmitoylation site. H8 holds a crucial position towards the intracellular side, where G-protein coupling occurs, and was shown to actively promote G-protein binding as well as stabilizing conformational states of the GPCR (Bruno et al., 2012; Palczewski et al., 2000). Importantly, the short stretch between TM7 and H8 and the proximal C-terminus beyond H8 were structurally shown to be in close contact with the G-protein and predicted to determine selective G protein activation in bovine rhodopsin (Tsai et al., 2019; Tsai et al., 2018).

To functionally define the involvement of these GPCR-G-protein contact regions, mutational studies were conducted, but the focus was on single residues and mutation most likely perturbed conformational and biophysical properties of the GPCR. Others compared the differences in active states of a GPCR known to bind to different G-proteins (Ma et al., 2020; Qiao et al., 2020). For example, b2AR binds primarily to the Gs-protein but can also stimulate the Gi/o pathway (Alegre et al., 2021; Xiao, 2001). These structures provided some insights into the residues involved in G-protein binding and potentially also G-protein selectivity, although the main differences were found to be different interaction modes of the GPCR with the G-protein (Alegre et al., 2021; Sandhu et al., 2019).

In this study, we functionally explored the involvement of the proximal C-terminal domain in G-protein selectivity. We employed engineered chimeric Opto-GPCRs, proteins based on an opsin GPCR with specific intracellular domains exchanged by those of a selected ligand-gated target receptor (Kleinlogel, 2016). Pioneering studies were performed with bovine rhodopsin (bRhod), where all intracellular domains were swapped by analogues of ligand-gated target receptors and the chimeras were tested functionally for their G-protein binding selectivity (Kim et al., 2005; Yamashita et al., 2001). However, as shown in recent work, engineering all intracellular domains results in poorer expression and functionality in in vivo experiments (Kralik et al., 2022). Thus, the second aim of this study was to determine which intracellular domains were crucial to replace to induce the desired downstream signaling. For this, we engineered a systematic assortment of chimeras between the structurally and functionally best known receptors, namely bovine rhodopsin (Gt and Gi selective (Terakita et al., 2002)) and the beta2-adrenoceptor (b2AR) (primarily Gs, secondarily Gi selective (Hasseldine et al., 2003; Rasmussen, DeVree, et al., 2011; Xiao, 2001)), as well as a set of chimeras between the not yet crystallized melanopsin (mainly Gq, partially Gi/o selective (Bailes & Lucas, 2013)) and mGluR6 (mainly Go selective, minimally Gi selective (Tian & Kammermeier, 2006)), two proteins with potentially different G-protein activation mechanisms (Seven et al., 2021; Spoida et al., 2016; Valdez-Lopez et al., 2020). We found that exchanging the proximal C-terminus was simultaneously crucial and sufficient to shift G-protein tropism in the melanopsin-mGluR6 chimera, whereas it was the known TM5-IL3-TM6 domain in the bRhod-b2AR chimera that induced the desired signaling.

## 2 Material & Methods

### 2.1 Sequences and cloning

The following GenBank sequences were used to engineer the opsin-target receptor chimeras: human melanopsin (AF147788.1), mGluR6 (U82083.1), bovine rhodopsin (AH001149.2), and beta2-adrenergic receptors (AY136741.1). Additionally, mouse melanopsin (AF147789.1) was used.

All chimeric opsin versions were designed as described in the main text, first cloned in pIRES_opsin_TurboFP635 plasmids and then modified to contain the fluorescent proteins mKate or mScarlet. To create the plasmids of the melanopsin-mGluR6 (pIRES_melanopsin-mGluR6_TurboFP635) and bRhod-b2AR chimeras (pIRES_rhodopsin-b2AR_TurboFP635), the respective domains of the intracellular loops and the C-terminus from mGluR6 or b2AR, respectively, were synthesized as primer overhangs and introduced using overlap extension PCR with EcoRI and BamHI restriction sites.

In the modified variants containing the fluorescent proteins, the Kir2.1 trafficking sequence (TS; KSRITSEGEYIPLDQIDINV) was inserted, as well as the 1D4 sequence from the C-terminus of bovine rhodopsin (TETSQVAPA), to create the final chimeras in pIRES_hMelanopsin-mGluR6_TS-1D4-mScarlet and pIRES_bRhod-b2AR_TS-1D4-mKate, respectively. All constructs were verified by Sanger sequencing (MicroSynth).

### 2.2 Cell Culture

HEK293 wildtype cells were cultured in Dulbecco’s modified Eagle medium (DMEM) in an incubator at 37° C with 5% CO_2_ atmosphere. DMEM was supplemented with 10% fetal calf serum (FCS; Seraglob, S70500), 1X glutamine (Seraglob, K8701), 1% penicillin/streptomycin (Sigma, P0781). Cells were seeded in 24-or 96-well plates (Greiner) at a density of 40’000 or 15’000 cells per well, respectively, and transiently transfected after 24 hours using the TransIT®LI 1 reagent (MirusBio, MIR2300) according to the manufacturer’s instructions. Briefly, the DNA and the transfection reagent were mixed in OptiMem medium (Sigma, 31985062) and incubated at room temperature for 20 minutes before transferring it to the cells. For 24-well plates, 0.5µg of plasmid DNA and 1.5µl of reagent was mixed in 50µl of OptiMem. For 96-well plates, 0.1µg of plasmid and 0.3µl of Mirus was mixed in 10µl of OptiMem. After transfection, all further steps were carried out under dim red light.

### 2.3 Plate Reader Assays

To test for specificity, we performed a variety of plate reader assays, including the aequorin assay to test for Gq-coupling, the GloSensor assay to test for Gi/o- and Gs-coupling, as well as the GsX assay to specify Gi/o- and Gq-coupling more precisely.

#### 2.3.1 Aequorin Ca2+ assay

To test for Gq-selectivity, we performed a standard Ca^2+^ assay using aequorin, a calcium-sensitive bioluminescent reporter protein that is used extensively in GPCR assays as a calcium indicator. The measured luminescence is directly proportional to the Ca^2+^ concentration. For transfection, cells were transiently transfected with a 2:1 ratio of GPCR chimera plasmid and reporter DNA. Cells were incubated in the transfection reagent for 3-4 hours and then supplemented by complete medium containing 1µg/ml doxycyclin and 1µM 9-cis retinal. The following day, the cells were incubated for one hour in the dark at room temperature in phenol-free Leibovitz (L-15) medium containing penicillin/streptomycin, 10% FCS, and L-glutamine (Gibco), 1µM 9-cis retinal and 10uM coelenterazine-h. For the measurements, 100 µl of cell suspension was added to a single well of a white 96-well plate. Luminescence was measured using an Infinite 200Pro Tecan plate reader (Männedorf, Switzerland). Raw luminescence was measured every 15 seconds. After letting the cells adapt for one minute and measuring baseline luminescence for six cycles, the plate was ejected, the cells were subjected to the stimulus, and recording resumed for a minimum of 15 additional cycles. Cells were stimulated by a single light flash. Data was collected in i-control (Tecan) from three replicates for each construct and condition during each assay.

#### 2.3.2 GloSensor cAMP assay

To test for Gs and Gi signaling, we performed a standard cAMP assay using GloSensor, a bioluminescent cAMP reporter. Cells were seeded at a density of 15’000 the day before or at 40’000 the day of transfection in solid white 96-well plates and co-transfected overnight or for at least 6 hours as above with a 2:1 ratio of GPCR plasmid and GloSensor reporter DNA. Cells were incubated at 37° and 5% CO2 in the transfection reagent for 1-2 hours before supplementing with medium containing 1µg/ml doxycyclin and 1µM 9-cis retinal and, where appropriate, 100ng pertussis toxin. Pertussis toxin was added to inhibit endogenous Gi signaling. The following day, 1-2 hours before taking the measurements, the cells were incubated at room temperature in the dark in phenol-red free Leibovitz media containing FBS, penicillin/streptomycin and L-glutamine, with 1µM 9-cis retinal and 4mM beetle luciferin. Beetle luciferin potassium salt (Promega) was reconstituted in 10mM HEPES with a p.H. of 6.9.

For the Gs second messenger assay, a baseline luminescence was measured for every 15 seconds for 12 cycles, then the plate was ejected, and the cells were subjected to the stimulus. For the Gi assay, 50µl of 3µM forskolin was added to the cells directly prior to beginning the assay to raise cAMP levels before exposing the cells to the stimulus. Here, raw luminescence was measured every 15 seconds for 60 cycles to allow for cAMP levels to rise high enough in order for proper measurement of the cAMP decrease following the light stimulus. Cells were stimulated with a single 470 nm light flash from a custom-built LED array. Data was collected from three replicates for each construct and condition during each assay. To evaluate the Gs to Gi signaling ratio for bRhod-b2AR chimeras, pertussis toxin (PTX) was added to the Gs assay, blocking Gi repsonses. Subtracting the Gs response during PTX block from the Gs response without PTX was used as a proxy for the pure Gi response.

#### 2.3.3 GsX assay

The GsX assay was established according to the protocol by Ballister and R. Lucas (Ballister et al., 2018). GsX plasmids (AddGene) are Gs subunit chimeras that transform the coupling of receptors to specific G proteins into increases in cAMP levels. Therefore, this assay can be used to test a greater variety of G protein coupling more precisely within one assay. Since the C-termini of the G proteins are known to be crucial in coupling selectivity, chimeric Gs subunits were designed by the Lucas group, in which the 13 most distal residues are replaced with different Galpha C-terminal sequences. Consequently, a GPCR that hypothetically couples to Gi, Gt and G15 would activate the GsX chimeras with a Gi, Gt or G15 tail (termed Gsi, Gst, and Gs15), leading to downstream Gs signaling and an increase in cAMP levels. The assays were executed analogously to the GloSensor cAMP assays described above. For transfection, cells were transfected with a 1:2 ratio of GloSensor reporter and plasmid DNA. Additionally, 50 ng of GsX chimera DNA was added to the transfection reagent, resulting in a final ratio of 10:5:1 of plasmid, reporter and GsX DNA. We found the 10:1 ratio of plasmid:GsX to be optimal when tested with human rhodopsin and Gsi (see supplementary Fig. S2).

### 2.4 Immunofluorescence

For immunohistochemistry, cells were transiently transfected in 24-well plates at a density of 40’000 cells as described above. Cells were then seeded onto coated Superfrost™ Plus Adhesion Microscope slides (Epredia, J1800AMNZ) and incubated at 37° overnight.

Sections were analyzed either under a Zeiss inverted microscope, equipped with Axiocam 712 mono-camera and ZEISS-Blue software, or under an ANA_Zeiss_LSM 880 confocal microscope (equipment supported by the Microscopy Imaging Center (MIC), University of Bern, Switzerland). Images were then processed and analyzed using ImageJ. The fluorescence intensity was measured across the cell membrane by drawing a line across the membrane and plotting the intensity profile in ImageJ, and was consecutively normalized in Excel.

### 2.5 Data processing and statistics

For the second messenger plate reader assays, raw luminescence was first normalized by dividing by the measured mKate2 fluorescence of each well, normalizing for the number of cells and subtracting the baseline luminescence signal. In the standard second messenger assays, the luminescence signal was normalized to baseline and then normalized to the wildtype opsin for the Gs and Gq assays, which was set to 1. For the Gi assays, luminescence was normalized to the maximum fold decrease in cAMP levels and then analogously normalized to the wildtype opsin. This is shown as the normalized luminescence in the time-course graphs in Suppl. Figs. 3,4,5 and 7. For the GsX assays, the maximum fold increase in the cAMP level was calculated and normalized to baseline. The wildtype opsin was analogously set to 1. In general, the analyses were performed on averaged, pooled data from the individual replicates. Given N values represent the number of triplets tested.

In the standard cAMP and Ca2+ second messenger assays, Gi activation efficacy was calculated by dividing maximum Gi signaling by maximum Gq signaling. G-protein signaling efficacy was calculated by adding the maximum Gi- and Gq responses and dividing by two. In the GsX assay, the relative activation efficacies of the different G-protein subtypes were calculated by dividing the maximum response of each GsX chimera by the sum of the maximum responses of all chimeras tested. The Gi activation efficacy was calculated by dividing the sum of Gsi, Gso and Gst responses by the sum of the Gsq and Gs15 responses. The G-protein signaling efficacy was calculated by dividing the sum of all maximum responses by 5.

All statistical analyses were performed either by Excel or using Graphpad Prism 9 software. In the figures the different levels of significance are indicated by * for p<0.05, ** for p<0.01, *** if p<0.001, and **** if p<0.0001. Average values are indicated with ± standard deviation.

### 2.6 Structural AlphaFold2 models

Alphafold-predicted structural models of our chimeras were calculated with ColabFold (Jumper et al., 2021; Mirdita et al., 2022) under default settings, alone and complexed with their G-proteins, with three recycling iterations and using MMseqs2 to search environmental and UniRef sequences. The protein structures were visualized, and images were captured using UCSF ChimeraX (Pettersen et al., 2021).

## 3 Results

### 3.1 Bovine rhodopsin – beta2 adrenoceptor chimeras (bRhod-b2ARs)

#### 3.1.1 Design and Engineering

We started by exploring chimeras between bRhod (GenBank accession number: AH001149.2) and b2AR (GenBank accession number: AY136741.1), since (1) the proximal C-terminus of bRhod was hypothesized to be involved in G-protein selectivity (Tsai et al., 2019), (2) abundant structural data is available, (3) both belong to Class A GPCRs facilitating sequence alignment (Kleinlogel, 2016) and (4) since functional rhodopsin chimeras with all intracellular domains exchanged have previously been successfully engineered (Airan et al., 2009; Kim et al., 2005; Tichy et al., 2022; Yamashita et al., 2001). Using the available structures as templates, we successively replaced IL2, IL3 and the C-terminus of rhodopsin by homologous domains of b2AR (Fig. 1A,B; all sequences are provided in Suppl. Data S1; an overview over all chimeras is given in Suppl. Table S1). In addition, the known G-protein contacts of bRhod and b2AR (Suppl. Fig. S3 A,B) were considered when defining the chimeric exchange points. In line with previous work (Airan et al., 2009; Kim et al., 2005), we included the intracellular ends of the TM5 and TM6 in the IL3 replacements as structural research has shown these regions to include important G-protein contact sites (Suppl. Fig. S3 A,B) (Rasmussen, DeVree, et al., 2011; Zhou et al., 2019).

**Figure 1:**
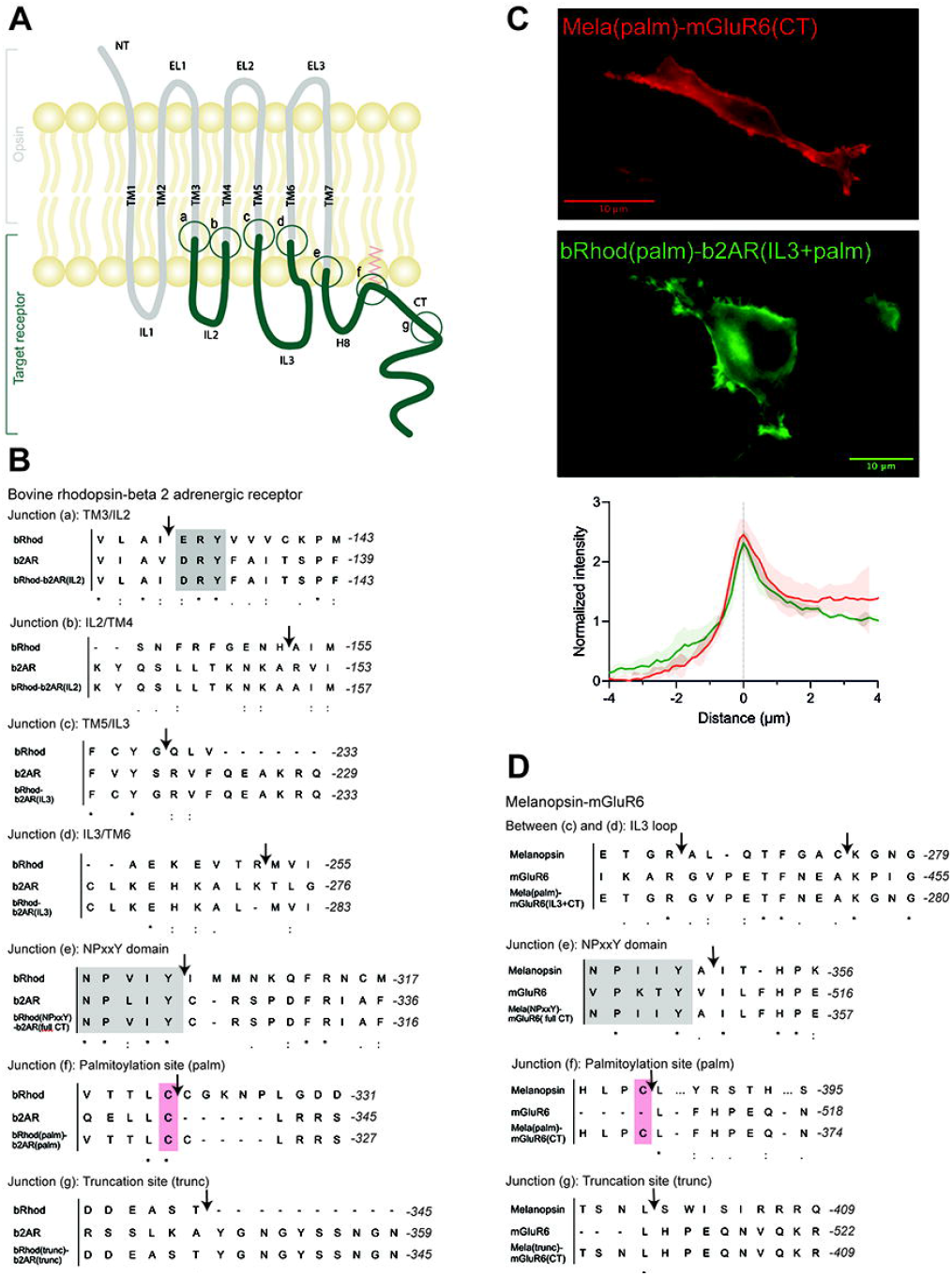
OptoGPCR engineering and expression in HEK293 cells. (A) Schematic representation of an Opto-GPCR chimera. The opsin domains are drawn in grey, the domains of the ligand-gated target receptor in dark green. Sites A-G highlight the chimeric recombination sites, detailed in panels B and D. (B,D) Sequence alignments of bovine rhodopsin with the beta-2 adrenergic receptor (B) and melanopsin with the mGluR6 receptor (D) around the chimeric fusion sites shown in (A). Conserved reference sites are highlighted in grey (NPxxY) and pink (palmitoylated cysteine). (C) Example photomicrographs and cross-sectional expression profiles of HEK293 cells expressing Mela(palm)-mGluR6(CT)-mKate2 and bRhod(palm)-b2AR(IL3+CT)-mKate2 fusion proteins, nicely trafficked to the cell membrane. Please refer to Suppl. Fig. S4 for all other chimeras.

The aim of the chimera design was to adapt G-protein specificity from that of bRhod to that of b2AR while maintaining functionality of the receptor. Since we were mainly interested in the functional contributions of the proximal C-terminus (from helix 8 to AA335 in bRhod), which was recently hypothesized from structure of bRhod (Tsai et al., 2019), we created three bRhod-b2AR chimeras that only contained b2AR domains within the C-terminus: (1) bRhod(trunc)-b2AR(CT), which retained rhodopsin’s C-terminus up to AA335, reported to modulate the signaling kinetics (Herrmann et al., 2006; Phillips & Cerione, 1994) and Gi-binding in rhodopsin (Tsai et al., 2019), (2) bRhod(palm)-b2AR(CT), retaining the palmitoylation site of rhodopsin at the end of helix 8 (H8) that anchors H8 into the cell membrane and mediates conformational stability of the receptor (Palczewski et al., 2000) and (3) bRhod(NPxxY)-b2AR(CT), where the whole C-terminus from the NPxxY motif at the distal end of TM7 was replaced. For a graphical view of the chimeric exchange sites refer to figure 1A-C and supplementary figure S3A,B.

All chimeric constructs were C-terminally tagged with the fluorescent mKate2 protein and cloned behind a CMV promoter for transient transfection of HEK293 cells. Fluorescent microscopy for mKate2 confirmed expression and correct membrane localization (Fig. 1C, Suppl. Fig. S4).

#### 3.1.2. Shifting G-protein selectivity from bRhod (Gi) to that of b2AR (Gs)

To test which chimeric design maximally shifts the response from the physiological Gi response of bRhod (Fig. 2A, Suppl. Fig. S5) to the Gs response of b2AR (Fig. 2B, Suppl. Fig. S5), we performed a standard bioluminescent cAMP plate reader assay in HEK293 cells (Bailes et al., 2012) to quantify the Gi-mediated decrease vs. the Gs-mediated increase in intracellular cAMP after light stimulation. As opposed to the findings by Tsai and colleagues, neither exchanging the full C-terminus [bRhod(NPxxY)-b2AR(CT)] (Tsai et al., 2018) nor exchanging the proximal C-terminus after H8 [bRhod(palm)-b2AR(CT)] (Tsai et al., 2019) in rhodopsin by that of b2AR changed G-protein tropism (Fig. 2A,B) from Gi to Gs, leading to Gs-activation (Fig. 2C). However, both C-terminal replacements decreased Gi-protein signaling (p=0.0016, p=0.0026, Fig. 2A). As expected, also replacing the full distal C-terminus beyond AA335 of bRhod [bRhod(trunc)-b2AR(CT)] did not change G-protein tropism (p=0.0053).

**Figure 2:**
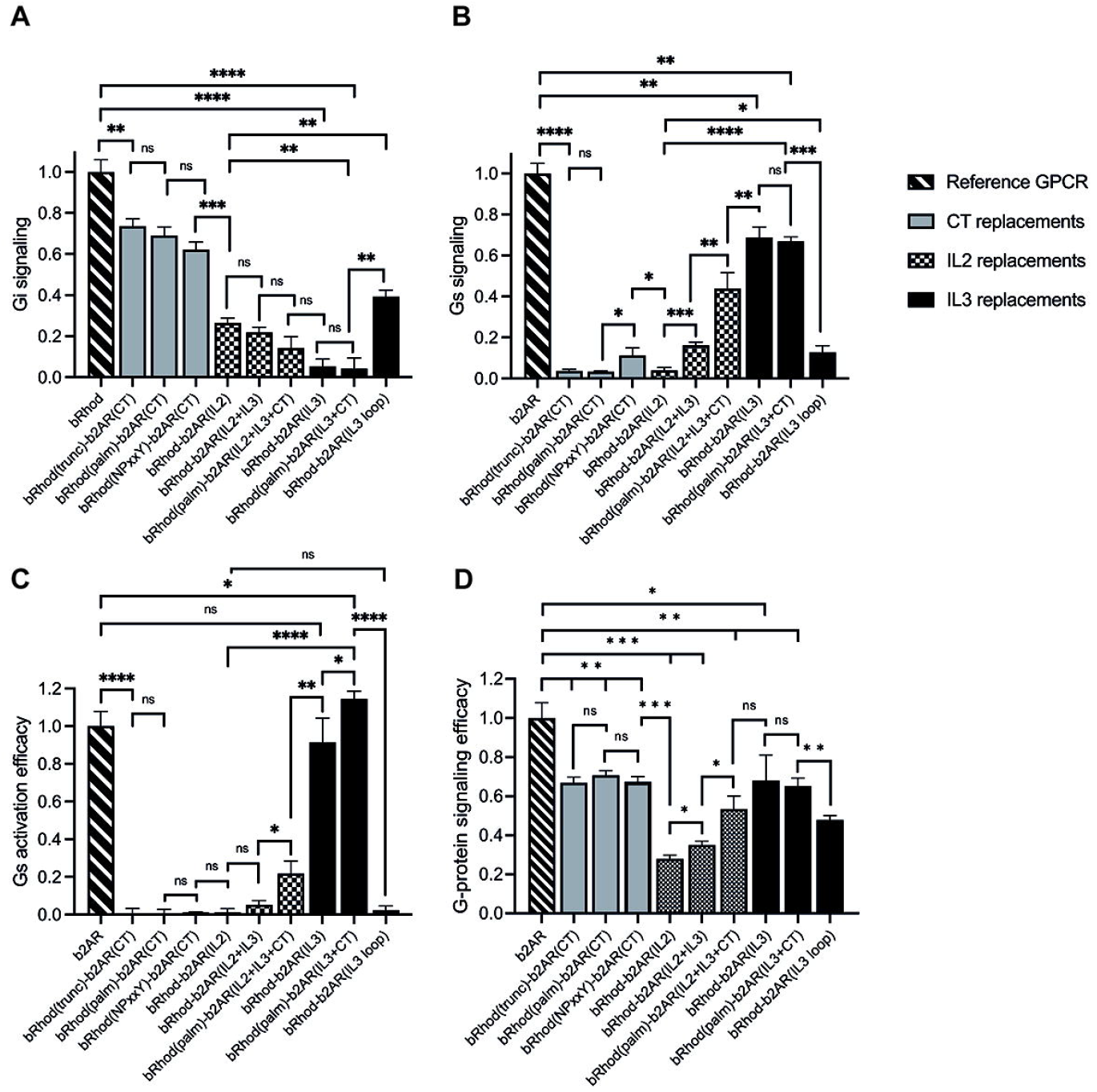
Shift from Gi to Gs activation of bRhod-b2AR chimeras determined with the GloSensor second messenger assay. (A,B) Normalized cAMP decrease indicative of Gi activation (A) and normalized increase indicative of Gs activation (B). The reporter luminescence values of the chimeras were normalized to WT bRhod (A) and WT b2AR (B) values, respectively (N=9). IL3 replacements, without and with the C-terminus additionally replaced at the palmitoylation site shifted bRhod-b2AR chimera signalling clearly to Gs. (C) Gs activation efficacy was calculated by dividing max. Gs value by the max. Gi value for each opsin variant. WT b2AR was normalized to 1, all other variants were normalized to WT b2AR (N=3). bRhod-b2AR(IL3) activated Gs as efficiently as WT b2AR, bRhod(palm)-b2AR(IL3+CT) significantly exceeded WT b2AR activity. (D) Overall G-protein signaling efficacy was calculated by dividing the sum of Gi and Gs activation by 2. Light stimulus: 5×10^16^ photons/cm^2^/s. Raw data traces can be found in Suppl. Fig. S6.

However, exchanging the full IL3, including the TM5 and TM6 helical cytoplasmic domains almost completely shifted G-protein selectivity to Gs (p<0.0001, Fig. 2, Suppl. Fig. S6), confirming existing literature that IL3 is the major mediator of G-protein selectivity (Flock et al., 2017; Rasmussen, DeVree, et al., 2011). Additionally exchanging the proximal C-terminus in bRhod(palm)-b2AR(IL3+CT) did not enhance Gs tropism (Fig. 2B), but significantly increased Gs activation (Fig. 2C; p=0.0423), corroborating a synergistic interaction of the proximal C-terminus with IL3 also in b2AR, as proposed by Tsai and colleagues for rhodopsin (Tsai et al., 2019). However, exchanging only the hypervariable loop region of IL3 was not as effective at changing G-protein tropism and reduced G-protein activation efficacy. This confirms the findings by the Kobilka group (Rasmussen, Choi, et al., 2011; Rasmussen, DeVree, et al., 2011) that the distal part of TM5 and the proximal part of TM6, extending well into the cytoplasmic space and comprising multiple G-protein contacts (Suppl. Fig. S3) actively support G-protein selection. Exchanging only the TM5 and TM6 cytoplasmic helical regions killed, as expected, the function of the chimera (Suppl. Fig. S6) (Deupi & Kobilka, 2007).

Finally, since IL2 is also in close contact with the G-protein (Rasmussen, DeVree, et al., 2011), we additionally created three chimeras in which we exchanged IL2. IL2 exchange alone did not adapt tropism to Gs and significantly reduced G-protein signaling (Fig. 2B,D); p<0.0001). However, if the IL2 was exchanged together with IL3, tropism shifted to Gs (Fig. 2B; p=0.0005) and G-protein signaling was significantly improved (Fig. 2D; p=0.0136). Additional replacement of the C-terminus after the palmytoilation site further increased Gs tropism (p=0.0036) and G-protein signaling (Fig. 2D; p=0.0102). Although not highly significant, these results may point towards a synergistic role of IL2, IL3 and the proximal C-terminus in G-protein selectivity, however, the Gs tropism and the Gs activation efficacy of bRhod-b2AR(IL2+IL3+palm) was significantly decreased compared to bRhod-b2AR(IL3) (Fig. 2B,C, p<0.0001 and p=0.0038), suggesting that too much engineering may impact functioning of the bRhod-b2AR receptor. Conclusively, IL3 appears to dictate G-protein selectivity in the bRhod-b2AR chimera.

### 3.2. Melanopsin-mGluR6 chimeras

#### 3.2.1 Design and engineering

Based on the sequences of melanopsin (GenBank accession number: AF147788) and mGluR6 (GenBank accession number: NM_000843.4), we designed chimeric Mela-mGluR6 variants, wherein different parts of the C-terminus – analogue to the bRhod-b2AR chimeras - were replaced without and together with IL3 of the corresponding regions of mGluR6 (Fig. 1D, sequences provided in Suppl. Data S1; chimera overview provided in Suppl. Table S1). Chimeric design of Mela-mGluR6 variants is complicated by the fact that melanopsin belongs to the class A GPCRs, whereas mGluR6 belongs to class C GPCRs, showing marked differences in structure and sequence ((Pin et al., 2003), Fig. 1D, Suppl. Fig. S3). Nevertheless, the well-preserved domains, including the E/DRY motif, the ionic lock and the NPxxY domain, can be used as reference points for sequence alignment (Kleinlogel, 2016), Suppl. Fig. S3). However, alignment of the C-terminal sequence is difficult, since mGluRs possess neither a palmitoylation site, nor an ordered H8 (Bruno et al., 2012). Further, melanopsin and other Class A GPCRs possess a long IL3 and a short IL2, whereas this arrangement is swapped in mGluR6 and other Class C GPCRs, with a short IL3 and a long IL2. We were therefore forced to only replace the hypervariable IL3 loop region (Matos-Cruz et al., 2011) of melanopsin with the short IL3 of mGluR6 (Fig. 1D, Suppl. Figs. 3,8).

Since the most effective bRhod-b2AR chimera in terms of Gs activation was the combined IL3+CT exchange variant, we combined the three different C-terminal replacements in the Mela-mGluR6 chimeras directly with IL3 loop replacements (Fig. 1A,D, Suppl. Fig. S3): (1) a Mela(trunc)-mGluR6(IL3+CT) chimera, where the C-terminus was exchanged at AA400 of melanopsin, comprising the analogous part of the proximal C-terminus of melanopsin reported to bind to the G-protein in bovine rhodopsin (Tsai et al., 2019), (2) a Mela(palm)-mGluR6(IL3+CT) chimera, where the C-terminus was exchanged just distal to H8 of melanopsin after the palmitoylation site at AA367 and (3) a fully truncated Mela(NPxxY)-mGluR6(IL3+CT) variant where the whole C-terminus of melanopsin distal to TM7 was replaced by that of mGluR6. In all three chimeras, the whole mGluR6 C-terminus, starting just distal to the NPxxY (VPKTY in mGluR6) motif was included (Suppl. Figs. S1, S3). Since previous studies indicated that the C-terminus of mGluRs is involved in direct coupling with G-proteins (Rondard et al., 2011; Seven et al., 2021), we additionally designed a Mela(palm)-mGluR6(short CT) chimera that comprised a shorter mGluR6 C-terminus not including the proximal C-terminus of mGluR6 (Suppl. Fig. S2).

The chimeric constructs were C-terminally tagged with mKate2 and cloned behind a CMV promoter for transient transfection of HEK-293 cells. Fluorescent microscopy for mKate2 confirmed expression and correct membrane localization (Fig. 1C, Suppl. Fig. S4).

#### 3.2.2 Shifting G-protein selectivity from melanopsin (Gq) to that of mGluR6 (Gi)

We first compared the different C-terminal replacement melanopsin variants that also had the IL3 loop region exchanged in the standard cAMP and Ca2+ Glosensor bioluminescent assays (Suppl. Fig. S9) (Bailes & Lucas, 2013). As expected, none of the variants activated Gs (not shown), but Gq (indicative of melanopsin) and Gi (indicative predominantly for mGluR6) at different ratios (Fig. 3, Suppl. Fig. S9). Mela(trunc)-mGluR6(IL3+CT) still dominantly activated Gq (62.3 ± 8.2% Gq, 37.7 ± 1.5% Gi), similar to wildtype melanopsin (76.6 ± 2.3% Gq, 23.4 ± 0.4% Gi), albeit with a significant increase in Gi tropism, which we attribute to the additional IL3 exchange and confirmed in the next set of experiments (see Fig. 3B). However, once the proximal part of the C-terminus of melanopsin was exchanged, the G-protein tropism switched to Gi [Mela(palm)-mGluR6(IL3+CT): Gi 69.2 ± 0.5%, Gq 30.8 ± 0.5%; Mela(NPxxY)-mGluR6(IL3+CT): Gi 70.1 ± 1.3%, Gq 29.9 ± 0.8%] and the Gi activation efficacy increased approximately 8-fold compared to WT melanopsin (Fig. 3B).

**Figure 3:**
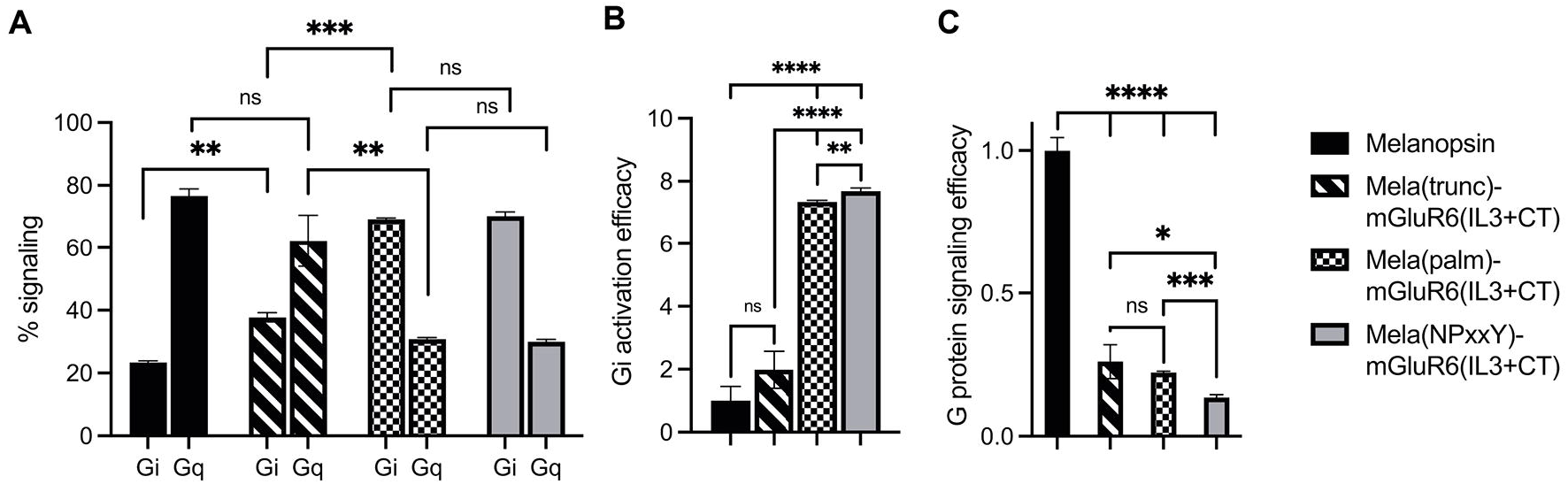
Shift from Gq to Gi activation of Mela-mGluR6 chimeras with combined IL3 and C-terminus replacements determined by the GsX second messenger assay. HEK293 cells transfected with the respective OptoGPCR plasmids, GloSensor reporter and the C-terminal mutated Gsq, Gs15, Gsi, Gso, or Gst G-proteins were subjected to a Gs cAMP assay. The maximum cAMP levels were divided by the sum of all values and normalized to 1 for each GsX protein. (A) The total Gq and Gi signaling was calculated by adding the max. cAMP value of Gsq and Gs15, and Gsi, Gso, and Gst, respectively, and dividing by the sum of all GsX responses to determine the percentage of each G-protein in total signalling. (B) The Gi activation efficacy was calculated to quantify the shift from Gq to Gi signaling. This was calculated by dividing the summed max. Gi values by the summed max. Gq value for each opsin variant. (C) The total G-protein signalling efficacy was calculated to determine the signaling functionality by calculating (Gi(total)+Gq(total))/2. WT melanopsin was normalized to 1, all other variants were normalized to WT melanopsin. Replacing the proximal C-terminus in melanopsin shifted G-protein specificity from Gq to Gi. However, disrupting the TM7-H8 domain of melanopsin in the Mela(NPxxY) variant significantly impacted the G-protein signalling efficacy. N=3; light stimulus: 470 nm, 5×10^16^ photons/cm^2^/s. Raw data traces can be found in Suppl. Fig. S9.

Evaluating the specific G-protein subtype activation profiles in a GsX assay (Ballister et al., 2018) (Suppl. Fig. S10) confirmed that the proximal C-terminus of melanopsin, from the NPxxY motif to AA400, dominantly determines G-protein selectivity in the Mela-mGluR6 chimera (Fig. 3, Suppl Fig. S10). Interestingly, the increase in Gi-protein tropism affected all three Gi-protein subtypes (Gi, Go and Gt) equally, hinting towards the fact that mGluR6 may not only activate Go and Gi (Tian & Kammermeier, 2006), but also Gt. However, the G-protein signaling efficacy of the Mela(NPxxY)-mGluR6(IL3+CT) chimera was significantly reduced compared to the Mela(palm)-mGluR6(IL3+CT) chimera (Fig. 3C; p<0.0001). We hypothesize that the lack of the amphipathic H8 in Mela(NPxxY)-mGluR6(IL3+CT) may destabilize conformational states and reduce G-protein activation upon light stimulation, an effect previously described for truncated rhodopsin (Krishna et al., 2002).

We went on to test another set of chimeras devoid of the IL3 loop replacement to investigate the explicit role of the proximal C-terminus in G-protein tropism. The results corroborated our findings above: while the Mela(trunc)-mGluR6(CT) version did not shift G-protein tropism, the Mela(palm)-mGluR6(CT) chimera shifted G-protein tropism from Gq to Gi (72.2 ± 0.2% Gi, 27.8 ± 1.1% Gq) almost identical to the Mela(palm)-mGluR6(IL3+CT) chimera (Fig. 4A). Consequently, the main determinant of G-protein specificity in melanopsin is the proximal C-terminus, with a subordinate role of IL3. It should also be noted that the Mela(palm)-mGluR6(CT) was significantly more effective in activating the G-proteins compared to the Mela(palm)-mGluR6(IL3+CT), additionally comprising the IL3 of mGluR6 (Figs 3C & 4B).

**Figure 4:**
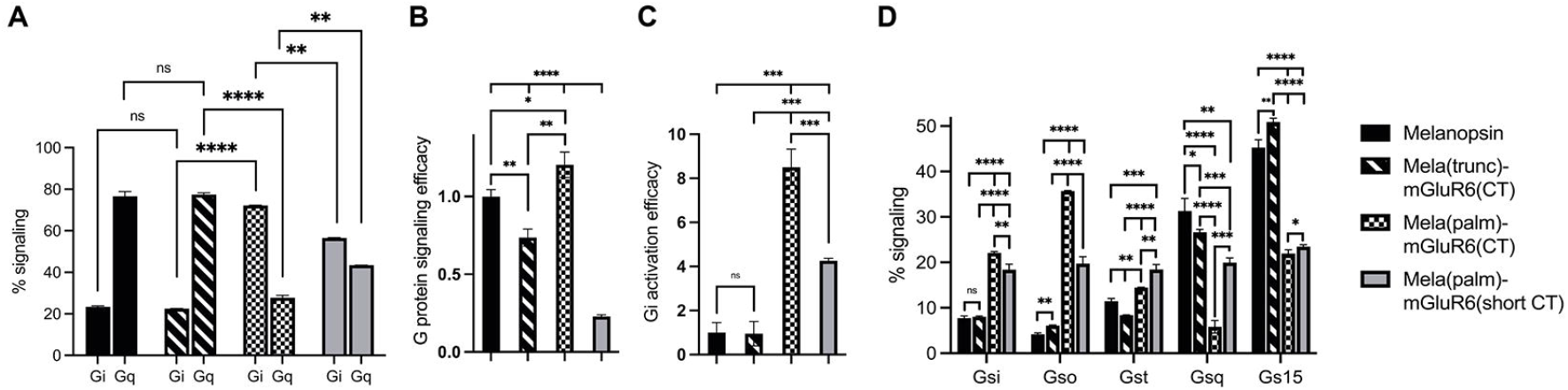
The role of the proximal C-terminus in Mela-mGluR6 chimeras in shifting tropism from Gq to Gi determined by the GsX second messenger assay. (A) The proximal C-terminus is responsible for G protein selectivity in the Mela-mGluR6 chimera and the analogue region steers G-protein selectivity also in mGluR6. The percentual proportion of each G-protein subfamily in total Gq and Gi signaling (100%) was calculated by dividing the max. cAMP response of each GsX-protein by the total sum of all max. GsX values. (B) The total G-protein signaling efficacy was determined by calculating (Gi(total)+Gq(total))/2. This was performed to quantify the functionality of the chimeric constructs. (C) The Gi activation efficacy ratio was calculated by dividing the summed mx. Gi values by the summed max. Gq values indicating the shift from Gq to Gi signaling. WT melanopsin was normalized to 1, all other variants were normalized to WT melanopsin. (D) The percentage of each G-protein in total signaling (100%) was calculated by dividing the max. cAMP response of each GsX-protein by the total sum of all GsX-proteins resulting in a fingerprint of the Gi and Gq signaling profiles of these variants. N=3; light stimulus: array of 3×8 LEDs, 470 nm, 5×10^16^ photons/cm^2^/s.

The seemingly mechanical difference of the Mela-mGluR6 chimera to the bRhod-b2AR chimera is remarkable. To visualize differences in the arrangement of the proximal C-terminus and IL3 in relation to the G-protein in the bRhod(palm)-b2AR(IL3+CT) and the Mela(palm)-mGluR6(CT) chimeras, we created structural AlphaFold models (Fig. 5A). Since no crystal structures are available for melanopsin or mGluR6, we modeled melanopsin on bovine rhodopsin (PDB: 6QNO) and mGluR6 on mGluR2 in complex with Gi (PDB: 7MTS). The bRhod-b2AR model shows proximity of the proximal C-terminus (residues AA375-AA382) to the Gbeta subunit as described previously (Tsai et al., 2019). The proximal C-terminus of the Mela-mGluR6 chimera, however, shows a very different, much more extended and tighter interface with the G-protein, starting immediately distal to H8 and lying within a cleft between the Galpha and Gbeta subunits with opposing hydrophobic (yellow) and hydrophilic (blue) surfaces. This defined placement and tight interaction with both, not only the Gbeta, but in particular also the Galpha subunit, may be the structural determinant of its involvement in G-protein selectivity. On the other hand, the IL3 of bRhod is in much closer proximity to the Galpha subunit, as this is the case in the Mela-mGluR6 chimera, hinting towards the dominant role of IL3 in G-protein selectivity.

**Fig. 5.**
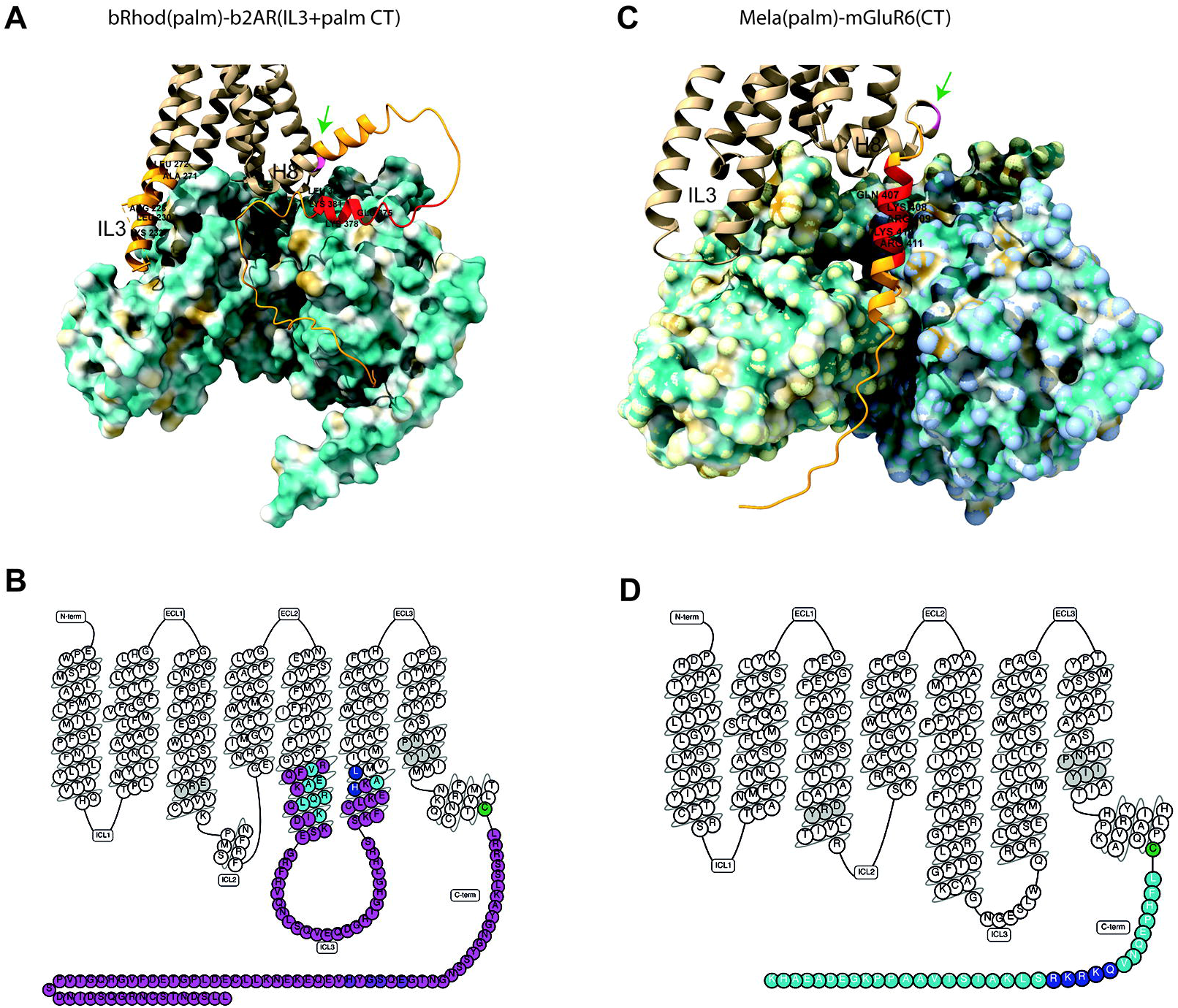
Structures of the bRhod(palm)-b2AR(IL3+CT) and Mela(palm)-mGluR6(CT) chimeras. (A,B) AlphaFold-predicted structural models of the chimeras with hydrophobicity surface maps of their respective G-proteins. (A) Structural model of the bRhod(palm)-b2AR(IL3+CT) chimera with the Gs-alpha and Gbeta subunits. Highlighted in light orange are the b2AR domains. The residues highlighted additionally in red are the analogues to the residues that form contacts with the Gbeta subunit in rhodopsin (Tsai et al., 2019). The palmitoylation site is indicated in pink (green arrow). (B) Structural model of the Mela(palm)-mGluR6(CT) construct with the Gi-alpha and Gbeta subunits. Highlighted in orange is the C-terminal domain of mGluR6. The residues highlighted in addition in red are analogous to the residues in mGluR2 that have been shown to form contacts with the Galpha subunit (Seven et al., 2021). (C,D) Snakeplots of the chimeras. (C) Snakeplot of the bRhod(palm)-b2AR(IL3+palm CT) construct. Shown in pink are the b2AR domains, which were inserted in bRhod. The residues that form contacts with the G-protein (light blue (Rasmussen, DeVree, et al., 2011) and dark blue (Zhou et al., 2019)) are highlighted, as well as the residues that are analogous to the residues that form contacts with the Gbeta subunit in rhodopsin (purple; (Tsai et al., 2019)). (D) Snakeplot of Mela(palm)-mGluR6(CT). Shown in light blue is the C-terminus of mGluR6, which was added following H8 and the palmitoylation site (green). The residues that are analogous to the residues that form contacts with the G-protein in mGluR2 (dark blue (Seven et al., 2021)) are highlighted. In grey are the highly conserved DRY and NPxxY motifs.

To investigate if the proximal C-terminus of mGluR6 is also involved in G-protein tropism, we next created an additional Mela(palm)-mGluR6(short CT) variant, where a truncated version of the mGluR6 C-terminus, commencing 3 AA downstream of the putative H8 considered devoid of G-protein interactions was employed (Rai et al., 2021) (Suppl. Fig. S3). Indeed, Gi activation was significantly reduced (Fig. 4C; p=0.0009) in this short CT variant in favor of Gq activation when compared to the full CT Mela(palm) version, although the overall tropism of the chimera remained shifted to Gi (Fig. 4A; 56.6 ± 0.2 % Gi, 43.4 ± 0.04% Gq). Defining the Gi-protein subtype specificity of Mela(palm)-mGluR6(short CT) and Mela(palm)-mGluR6(full CT) in more detail with the GsX assay clearly showed a shift predominantly to Go of mGluR6 (Tian & Kammermeier, 2006) only in the full CT version (Fig. 4D), proving that the proximal C-terminus is also the mediator of G-protein selectivity in mGluR6.

While we established the GsX assay, we found varying G-protein tropisms for murine and human melanopsin: human melanopsin had a significantly lower Gq specificity, but an elevated Gi specificity compared to murine melanopsin (Suppl. Fig. S11; murine: Gq 67.8 ± 0.8%, Gi 32.2 ± 1.07%; human: Gq 63.2 ± 0.4, Gi 36.8 ± 0.2). Interestingly, human melanopsin also possessed a significantly increased G15 *vs*. Gq tropism, which was not found in murine melanopsin (Suppl. Fig. 11). Intriguingly, when comparing the sequences of human and murine melanopsin, they deviated from each other exactly in the proximal C-terminus between H8 and the truncation site in Mela(trunc) (AA400), as well as the IL3, but not in the IL2 (Suppl. Fig. S11). This corroborates our findings that the proximal C-terminus and the IL3 influence G-protein selectivity in melanopsin.

Taken together, our results unravel the dominant role of the proximal C-terminus in G-protein selectivity in melanopsin and mGluR6.

## 4 Discussion

Selective coupling of GPCRs to specific G-protein subtypes is critical to transform signals from light, neurotransmitters, and drugs into intracellular responses throughout the body. In the past 30 years, the field has tried to understand the molecular and structural underpinnings in GPCRs responsible for G-protein selectivity, however, they remain largely elusive. The recent surge in resolved GPCR-G-protein structures, including structures in the active and inactive states, has expanded our understanding of G-protein recognition and GPCR-mediated signal transduction. However, since structural data only provide snapshots of particular GPCR-G-protein states and since the intracellular G-protein interacting domains on the GPCR (ILs and CT) are flexible, largely unordered domains, deferring functional involvement in G-protein selectivity remains a challenge (Deupi & Kobilka, 2007). Fab fragments (Rasmussen et al., 2007), mini G-proteins (Carpenter et al., 2016; Tsai et al., 2018) and nanobodies (Bertheleme et al., 2013) have been employed to immobilize these flexible domains to promote crystallization, which may, however, partially perturb dynamic movements of the GPCR and cast a distorted picture on GPCR/G-protein interactions. Opto-GPCRs present a powerful tool to add temporally longitudinal functional data of G-protein coupling signatures to the existing structural knowledge.

The first structure of the b2AR was obtained in the presence of a Fab fragment bound to IL3 (Rasmussen et al., 2007) and the first active state crystal structure was derived from rhodopsin (Palczewski et al., 2000). Most of the insights of GPCR-G-protein interactions have been derived from these two proteins and the residues described to be involved in G-protein contacts are similar (Suppl. Fig. S3) (Rasmussen, Choi, et al., 2011; Rasmussen, DeVree, et al., 2011; Tsai et al., 2018). With well-defined contacts to the G-protein (Flock et al., 2017; Kang et al., 2018; Rasmussen, DeVree, et al., 2011; Tsai et al., 2019; Zhou et al., 2019) and supporting functional data (Airan et al., 2009; Kim et al., 2005; Ma et al., 2020; Siuda et al., 2015), the IL3 is considered the main determinant of G-protein selectivity, at least in Class A GPCRs. Here, we were mainly interested in elucidating the potential involvement of the proximal C-terminus, which was also predicted to be involved in G-protein activation and selectivity in recent structural studies (Seven et al., 2021; Tsai et al., 2019). Tsai and colleagues showed the TM7/H8 joint of rhodopsin (Gao et al., 2019; Kang et al., 2018; Tsai et al., 2018) to AA(T/A335) (Tsai et al., 2019) to closely contact the G-protein and suggested a role in G-protein activation and potentially also selectivity. While we could not confirm a role in selectivity of the proximal C-terminus in our bRhod-b2AR chimeras, we found a significant decrease of Gi activation the more of the C-terminus of rhodopsin was replaced by that of b2AR, which corroborates the suggested importance of the proximal carboxyl terminus (from NPxxY to AA335) specifically in Gi-protein binding and activation. Our structural model of the bRhod(palm)-b2AR(palm+IL3) chimera visualizes the proximity of the proximal C-terminus to the Gbeta subunit, specifically of amino acids 375-382 (Fig. 5A), underscoring the supportive role of this domain in G-protein activation.

IL3 is one of the most diverse regions in GPCRs and is often structurally not completely resolved. For instance, b2AR has a much longer and extremely flexible IL3 compared to rhodopsin. IL3 exchange in our bRhod-b2AR chimera almost completely shifted G-protein selectivity to Gs. These findings agree with the common assumption that IL3 is the major mediator of G-protein specificity in bRhod and b2AR (Flock et al., 2017; Rasmussen, DeVree, et al., 2011). Exchanging only the hypervariable loop region of IL3 was not as effective as exchanging the entire intracellular IL3 domain including the intracellular TM5 and TM6 protrusions. This confirms the findings by the Kobilka group (Rasmussen, Choi, et al., 2011; Rasmussen, DeVree, et al., 2011) that the distal part of TM5 and the proximal part of TM6, extending into the cytoplasmic space and comprising multiple G-protein contact sites, actively support G-protein selection (Suppl. Fig. S3). However, only replacing TM5 and TM6 did not suffice to induce a Gs signal. Therefore, we conclude that the entire TM5-IL3-TM6 region is necessary for optimal Gs coupling of the bRhod-b2AR chimera. Exchanging the C-terminus in addition to IL3 in the bRhod-b2AR chimera (bRhod(palm)) had no significant effect on G-protein selectivity, but significantly increased Gs activation, confirming a supportive role of the proximal C-terminus in G-protein activation, probably by the hypothesized interactions with IL3 (Tsai et al., 2019).

Similarly, we did not find IL2 to promote Gs selectivity on its own, albeit shown to be in close contact to the G-protein in rhodopsin (Kang et al., 2018; Tsai et al., 2018), in b2AR (Rasmussen, DeVree, et al., 2011) and in our bRhod-b2R model (Fig. 5A). In fact, replacement of IL2 in the bRhod-b2AR chimera reduced the G-protein activation efficacy, suggesting that too much engineering in Opto-GPCRs may disturb the conformational states of the GPCR and/or hamper G-protein binding. A previous study on b1AR, a related receptor to b2AR also coupling to Gs, suggested that PIP2 lipid binding to the distal IL2 (cytoplasmic end of TM4) stabilizes the active state of b1AR by forming a PIP2 bridging interaction specifically to Gs (Yen et al., 2018). However, our bRhod-b2AR(IL2) chimera was still mainly sensitive to Gi. Potentially, the surrounding membrane environment may have affected PIP2 lipidation of IL2 leading to an underestimated Gs binding in our experiments.

While the results obtained with the bRhod-b2AR chimera are in good agreement with existing structural und mutational data (Kang et al., 2018; Rasmussen, DeVree, et al., 2011; Tsai et al., 2019; Zhou et al., 2019), we obtained novel results with the melanopsin-mGluR6 chimera. We discovered that the proximal C-termini of melanopsin (from the palmitoylation site at C367 to AA400) and of mGluR6 (AA846-AA864) act as main mediators of G-protein selectivity. It has to be noted, that neither the melanopsin nor the mGluR6 crystal structures are known.

The most effective variant to shift tropism from Gq to Gi/o was the Mela(palm)-mGluR6(CT) chimera. The fully truncated Mela(NPxxY)-mGluR6 variant was much less effective in G-protein activation, assumingly due to the lack of the amphipathic H8 shown to stabilize conformational states in rhodopsin, and potentially other opsins (Krishna et al., 2002). Valdez-Lopez and colleagues (Valdez-Lopez et al., 2020) suggested ionic interactions between the proximal melanopsin C-terminus (C367 - Y382) and IL3, specifically within the helical intracellular extensions of TM5 (R259, R262, R266, R277) and in the IL3 loop region (R280, Q281, W282, Q283, R284, L285), forming a stable conformation critical for initiating G-protein signaling. However, in our study, additionally replacing IL3 of melanopsin by that of mGluR6 significantly reduced the capacity of the chimera to activate the G-protein, hinting towards conformational disturbances. It has to be noted that IL3 exchange was not trivial in the melanopsin-mGluR6 chimera, since melanopsin belongs to Class A GPCRs, whereas mGluR6 belongs to Class C GPCRs. Melanopsin and other Class A GPCRs possess a long IL3 and a short IL2, whereas mGluR6 and other Class C GPCRs possess a short IL3 and a long IL2 (Pin et al., 2003). We were therefore forced to only replace the hypervariable IL3 loop region of melanopsin with the short IL3 of mGluR6, which potentially disturbed the suggested ionic network between IL3 and the proximal C-terminus. According to the sequence differences between human and murine melanopsin IL3, including the TM5 and TM6 extensions (Suppl. Fig. S11), for which we also found differences in G-protein selectivity, we suggest that Gi/o selectivity of the Mela(palm)-mGluR6(IL3+CT) chimera may be further enhanced by including the TM5 and TM6 protrusions into the IL3 replacement.

It is interesting that Valdez-Lopez also pointed towards this same proximal C-terminal region in Gq-coupled melanopsin, as residues on the proximal C-terminus are quite conserved across Gs and Gi proteins (Tsai et al., 2019), but not Gq proteins. Our structural model of the Mela(palm)-mGluR6(CT) chimera shows a very different arrangement of the C-terminus compared to the bRhod-b2AR chimera (Fig. 5). While in the bRhod-b2AR model, the proximal C-terminus contacts solely the Gbeta subunit, in the Mela(palm)-mGluR6(CT) model, the whole C-terminus is nested into a cleft between the Galpha and the Gbeta subunits. This difference in arrangement, and potentially the additional contact to the Galpha subunit, may explain the strong involvement of the proximal C-terminus in G-protein selectivity in the Mela(palm)-mGluR6(CT) chimera. The fact that the bRhod-b2AR chimera has a much closer contact of IL3 to the Galpha subunit corroborates that the contact to the Galpha subunit determines G-protein selectivity.

Already in the 1990s it was suggested from mGluR chimeras that the C-terminal segment located downstream of TM7 is necessary for specific G-protein activation (Pin et al., 1994). This was recently confirmed in mGluR2 by cryo-EM studies (Seven et al., 2021), which show a binding between the proximal C-terminus of mGluR2 and the Gi-protein. To study the involvement of the proximal C-terminus of mGluR6 in G-protein selectivity, we compared Mela(palm)-mGluR6(CT)-mGluR6 chimeras comprising the whole mGluR6 C-terminus (cut after the NPxxY motif) and a short C-terminus (Mela(palm)-mGluR6(short CT)), lacking the mGluR6 analog of the proximal C-terminus to melanopsin. Albeit definition of the proximal mGluR6 C-terminus is not trivial due to the lack of a palmitoylation site and an ordered H8 (Bruno et al., 2012) (see Suppl. Fig. S3), we could functionally confirm that the proximal C-terminus of mGluR6 (Q849-R856) comprising the analogue amino acid stretch mentioned by Seven and colleagues in mGluR2 (V826-S833) (Seven et al., 2021) (Suppl. Fig. 1), is an important mediator of G protein selectivity.

Considering Opto-GPCR variants as optogenetic tools, we showed in our study that in contrast to most previous work with Opto-GPCRs, in which the IL2, IL3 and the full C-terminus were replaced, only certain domains need to be replaced to induce the required shift in G-protein coupling without compromising the G-protein coupling efficacy. While the bRhod-b2AR variant was optimally designed by introducing the b2AR TM5-IL3-IL6 and C-terminus after the palmitoylation site in bRhod, a proximal C-terminus replacement of melanopsin by the whole C-terminus of the mGluR6 receptor, including the proximal C-terminus, sufficed to retarget to the Gi/o protein class. Overall, the Mela(palm)-mGluR6(CT) seems to be the favorable chimera for functional applications since it activates G-proteins with high efficacy and shows a strong tropism towards Gi/o (Kralik et al., 2022). Of course, since Cryo-EM studies on mGluR2 showed that the G-protein’s C-terminus is stabilized by a pocket formed by IL2 and the C-terminus (Seven et al., 2021), IL2 may also be involved in Gi/o-selectivity together with the proximal C-terminus of mGluR6, which was, however, not investigated in this study. Interestingly, Gi-protein subtype specificity (Gi, Go and Gt) was increased the more of the melanopsin C-terminus was replaced by that of mGluR6, hinting toward the fact that mGluR6 may not only activate Go and Gi (Tian & Kammermeier, 2006), but also Gt. This could be the reason why middle-wave cone opsin (Gaub et al., 2014) and rhodopsin (Cehajic-Kapetanovic et al., 2015), which both naturally bind to Gt (transducin), were efficient in activating ON-bipolar cells of the retina and restoring some vision. We also showed that bovine rhodopsin binds not only to Gt, but equally well to Gi and to a lesser degree (approx. 10%) to Go (Suppl. Fig. S5). This further explains why rhodopsin functioned well as an optogenetic tool in retinal-ON bipolar cells (Cehajic-Kapetanovic et al., 2015), since rhodopsin and mGluR6 both activate Gi, Go and Gt, just with different efficacies. Overall, it appears that GPCRs are far more promiscuous than generally believed.

Our study highlights the complexity and individuality of G-protein selectivity in GPCRs and with that the challenge to design Opto-GPCRs targeting a specific G-protein. From the limited results we generated with the bRhod-b2AR and Mela-mGluR6 chimeras, we suggest that GPCR interactions with the Galpha interface may trigger G-protein specificity, while interactions with the Gbeta protein support G-protein activation. From our results, no single blueprint can be created that would be applicable to all opsins and target receptors. Rather, there is a need to consider each opsin-target receptor pair separately to achieve efficient and optimized Opto-GPCR chimeric receptor function. To further corroborate these results and propose potential general schemes for different classes of GPCRs and for GPCRs that couple to different G-proteins, additional chimeras should be created and investigated. In the future, the gained knowledge will assist the design and functionality of Opto-GPCRs and drugs that promote specific signaling pathways and avoid unwanted side effects (Hauser et al., 2017).

## Author Contributions

SL performed and analyzed the experiments and wrote the first draft of the manuscript. SK supervised the project, designed the experiments, helped with the analysis and writing of the manuscript as well as provided funding for the project.

## Funding

This research was funded by grants from the Swiss National Science Foundation (310030E_188991) and 31003A_176065) to SK.

## Acknowledgments

We would like to thank Sabine Schneider for assistance in cloning, immunohistochemistry and the plate reader assays and Michiel van Wyk for the fruitful discussions about GPCRs and G-proteins.

## Manuscript contribution to the field

G-protein coupled receptors ubiquitously modulate physiology throughout our bodies. Therefore, they also present primary targets for pharmaceutical drugs. However, GPCR modulation is highly complex and despite recent major advances in available GPCR structures, the mechanism underlying selective G-protein binding, which mediates specific intracellular signalling remains still largely unknown. Understanding the mechanism of G-protein selectivity may in the future enable the design of drugs and optogenetic Opto-GPCRS that can activate specific intracellular pathways at will. In this study, we employed light-activatable Opto-GPCRs, in particular rhodopsin-beta2-adrenoceptor chimeras and melanopsin-mGluR6 chimeras, as tools to define the GPCR domains defining G-protein selectivity with a focus on the proximal C-terminus. We show that in melanopsin and mGluR6 the proximal C-terminus is the dominator of G-protein specificity, whereas it is the IL3 in bovine rhodopsin and beta2-adrenoceptor that define G-protein tropism. Collectively, this work adds knowledge to the GPCR domains mediating G-protein selectivity, ultimately paving the way to optogenetically elicited specific G-protein signalling on demand.

## Notes

### Competing Interest Statement

The authors have declared no competing interest.

## Literature

Airan, R. D., Thompson, K. R., Fenno, L. E., Bernstein, H., & Deisseroth, K. (2009). Temporally precise in vivo control of intracellular signalling. Nature, 458(7241), 1025–1029. https://doi.org/10.1038/nature07926

Alegre, K. O., Paknejad, N., Su, M., Lou, J. S., Huang, J., Jordan, K. D., Eng, E. T., Meyerson, J. R., Hite, R. K., & Huang, X. Y. (2021). Structural basis and mechanism of activation of two different families of G proteins by the same GPCR. Nat Struct Mol Biol, 28(11), 936–944. https://doi.org/10.1038/s41594-021-00679-2

Bailes, H. J., & Lucas, R. J. (2013). Human melanopsin forms a pigment maximally sensitive to blue light (lambdamax approximately 479 nm) supporting activation of G(q/11) and G(i/o) signalling cascades. Proc Biol Sci, 280(1759), 20122987. https://doi.org/10.1098/rspb.2012.2987

Bailes, H. J., Zhuang, L. Y., & Lucas, R. J. (2012). Reproducible and sustained regulation of Galphas signalling using a metazoan opsin as an optogenetic tool. PLoS One, 7(1), e30774. https://doi.org/10.1371/journal.pone.0030774

Ballister, E. R., Rodgers, J., Martial, F., & Lucas, R. J. (2018). A live cell assay of GPCR coupling allows identification of optogenetic tools for controlling Go and Gi signaling. BMC Biol, 16(1), 10. https://doi.org/10.1186/s12915-017-0475-2

Bertheleme, N., Chae, P. S., Singh, S., Mossakowska, D., Hann, M. M., Smith, K. J., Hubbard, J. A., Dowell, S. J., & Byrne, B. (2013). Unlocking the secrets of the gatekeeper: methods for stabilizing and crystallizing GPCRs. Biochim Biophys Acta, 1828(11), 2583–2591. https://doi.org/10.1016/j.bbamem.2013.07.013

Bruno, A., Costantino, G., de Fabritiis, G., Pastor, M., & Selent, J. (2012). Membrane-sensitive conformational states of helix 8 in the metabotropic Glu2 receptor, a class C GPCR. PLoS One, 7(8), e42023. https://doi.org/10.1371/journal.pone.0042023

Carpenter, B., Nehme, R., Warne, T., Leslie, A. G., & Tate, C. G. (2016). Structure of the adenosine A(2A) receptor bound to an engineered G protein. Nature, 536(7614), 104–107. https://doi.org/10.1038/nature18966

Cehajic-Kapetanovic, J., Eleftheriou, C., Allen, A. E., Milosavljevic, N., Pienaar, A., Bedford, R., Davis, K. E., Bishop, P. N., & Lucas, R. J. (2015). Restoration of Vision with Ectopic Expression of Human Rod Opsin. Curr Biol, 25(16), 2111–2122. https://doi.org/10.1016/j.cub.2015.07.029

Choe, H. W., Kim, Y. J., Park, J. H., Morizumi, T., Pai, E. F., Krauss, N., Hofmann, K. P., Scheerer, P., & Ernst, O. P. (2011). Crystal structure of metarhodopsin II. Nature, 471(7340), 651–655. https://doi.org/10.1038/nature09789

Deupi, X., & Kobilka, B. (2007). Activation of G protein-coupled receptors. Adv Protein Chem, 74, 137–166. https://doi.org/10.1016/S0065-3233(07)74004-4

Flock, T., Hauser, A. S., Lund, N., Gloriam, D. E., Balaji, S., & Babu, M. M. (2017). Selectivity determinants of GPCR-G-protein binding. Nature, 545(7654), 317–322. https://doi.org/10.1038/nature22070

Gao, Y., Hu, H., Ramachandran, S., Erickson, J. W., Cerione, R. A., & Skiniotis, G. (2019). Structures of the Rhodopsin-Transducin Complex: Insights into G-Protein Activation. Mol Cell, 75(4), 781–790 e783. https://doi.org/10.1016/j.molcel.2019.06.007

Gaub, B. M., Berry, M. H., Holt, A. E., Reiner, A., Kienzler, M. A., Dolgova, N., Nikonov, S., Aguirre, G. D., Beltran, W. A., Flannery, J. G., & Isacoff, E. Y. (2014). Restoration of visual function by expression of a light-gated mammalian ion channel in retinal ganglion cells or ON-bipolar cells. Proc Natl Acad Sci U S A, 111(51), E5574–5583. https://doi.org/10.1073/pnas.1414162111

Hasseldine, A. R., Harper, E. A., & Black, J. W. (2003). Cardiac-specific overexpression of human beta2 adrenoceptors in mice exposes coupling to both Gs and Gi proteins. Br J Pharmacol, 138(7), 1358–1366. https://doi.org/10.1038/sj.bjp.0705191

Hauser, A. S., Attwood, M. M., Rask-Andersen, M., Schioth, H. B., & Gloriam, D. E. (2017). Trends in GPCR drug discovery: new agents, targets and indications. Nat Rev Drug Discov, 16(12), 829–842. https://doi.org/10.1038/nrd.2017.178

Herrmann, R., Heck, M., Henklein, P., Kleuss, C., Wray, V., Hofmann, K. P., & Ernst, O. P. (2006). Rhodopsin-transducin coupling: role of the Galpha C-terminus in nucleotide exchange catalysis. Vision Res, 46(27), 4582–4593. https://doi.org/10.1016/j.visres.2006.07.027

Jumper, J., Evans, R., Pritzel, A., Green, T., Figurnov, M., Ronneberger, O., Tunyasuvunakool, K., Bates, R., Zidek, A., Potapenko, A., Bridgland, A., Meyer, C., Kohl, S. A. A., Ballard, A. J., Cowie, A., Romera-Paredes, B., Nikolov, S., Jain, R., Adler, J., … Hassabis, D. (2021). Highly accurate protein structure prediction with AlphaFold. Nature, 596(7873), 583–589. https://doi.org/10.1038/s41586-021-03819-2

Kang, Y., Kuybeda, O., de Waal, P. W., Mukherjee, S., Van Eps, N., Dutka, P., Zhou, X. E., Bartesaghi, A., Erramilli, S., Morizumi, T., Gu, X., Yin, Y., Liu, P., Jiang, Y., Meng, X., Zhao, G., Melcher, K., Ernst, O. P., Kossiakoff, A. A., … Xu, H. E. (2018). Cryo-EM structure of human rhodopsin bound to an inhibitory G protein. Nature, 558(7711), 553–558. https://doi.org/10.1038/s41586-018-0215-y

Kim, J. M., Hwa, J., Garriga, P., Reeves, P. J., RajBhandary, U. L., & Khorana, H. G. (2005). Lightdriven activation of beta 2-adrenergic receptor signaling by a chimeric rhodopsin containing the beta 2-adrenergic receptor cytoplasmic loops. Biochemistry, 44(7), 2284–2292. https://doi.org/10.1021/bi048328i

Kleinlogel, S. (2016). Optogenetic user’s guide to Opto-GPCRs. Front Biosci (Landmark Ed), 21(4), 794–805. https://doi.org/10.2741/4421

Kralik, J., Wyk, M. v., Stocker, N., & Kleinlogel, S. (2022). Bipolar cell targeted optogenetic gene therapy restores parallel retinal signaling and high-level vision in the degenerated retina. Commun Biol. https://doi.org/10.21203/rs.3.rs-999738/v1

Krishna, A. G., Menon, S. T., Terry, T. J., & Sakmar, T. P. (2002). Evidence that helix 8 of rhodopsin acts as a membrane-dependent conformational switch. Biochemistry, 41(26), 8298–8309. https://doi.org/10.1021/bi025534m

Lebon, G., Warne, T., Edwards, P. C., Bennett, K., Langmead, C. J., Leslie, A. G., & Tate, C. G. (2011). Agonist-bound adenosine A2A receptor structures reveal common features of GPCR activation. Nature, 474(7352), 521–525. https://doi.org/10.1038/nature10136

Ma, X., Hu, Y., Batebi, H., Heng, J., Xu, J., Liu, X., Niu, X., Li, H., Hildebrand, P. W., Jin, C., & Kobilka, B. K. (2020). Analysis of beta2AR-Gs and beta2AR-Gi complex formation by NMR spectroscopy. Proc Natl Acad Sci U S A, 117(37), 23096–23105. https://doi.org/10.1073/pnas.2009786117

Matos-Cruz, V., Blasic, J., Nickle, B., Robinson, P. R., Hattar, S., & Halpern, M. E. (2011). Unexpected diversity and photoperiod dependence of the zebrafish melanopsin system. PLoS One, 6(9), e25111. https://doi.org/10.1371/journal.pone.0025111

Milligan, G., & Kostenis, E. (2006). Heterotrimeric G-proteins: a short history. Br J Pharmacol, 147 Suppl 1, S46–55. https://doi.org/10.1038/sj.bjp.0706405

Mirdita, M., Schutze, K., Moriwaki, Y., Heo, L., Ovchinnikov, S., & Steinegger, M. (2022). ColabFold: making protein folding accessible to all. Nat Methods, 19(6), 679–682. https://doi.org/10.1038/s41592-022-01488-1

Neves, S. R., Ram, P. T., & Iyengar, R. (2002). G protein pathways. Science, 296(5573), 1636–1639. https://doi.org/10.1126/science.1071550

Oldham, W. M., & Hamm, H. E. (2008). Heterotrimeric G protein activation by G-protein-coupled receptors. Nat Rev Mol Cell Biol, 9(1), 60–71. https://doi.org/10.1038/nrm2299

Palczewski, K., Kumasaka, T., Hori, T., Behnke, C. A., Motoshima, H., Fox, B. A., Le Trong, I., Teller, D. C., Okada, T., Stenkamp, R. E., Yamamoto, M., & Miyano, M. (2000). Crystal structure of rhodopsin: A G protein-coupled receptor. Science, 289(5480), 739–745. https://doi.org/10.1126/science.289.5480.739

Pettersen, E. F., Goddard, T. D., Huang, C. C., Meng, E. C., Couch, G. S., Croll, T. I., Morris, J. H., & Ferrin, T. E. (2021). UCSF ChimeraX: Structure visualization for researchers, educators, and developers. Protein Sci, 30(1), 70–82. https://doi.org/10.1002/pro.3943

Phillips, W. J., & Cerione, R. A. (1994). A C-terminal peptide of bovine rhodopsin binds to the transducin alpha-subunit and facilitates its activation. Biochem J, 299 (Pt 2), 351–357. https://doi.org/10.1042/bj2990351

Pin, J.-P., Galvez, T., & Prézeau, L. (2003). Evolution, structure, and activation mechanism of family 3/C G-protein-coupled receptors. Pharmacology & Therapeutics, 98(3), 325–354. https://doi.org/10.1016/s0163-7258(03)00038-x

Pin, J. P., Joly, C., Heinemann, S. F., & Bockaert, J. (1994). Domains involved in the specificity of G protein activation in phospholipase C-coupled metabotropic glutamate receptors. EMBO J, 13(2), 342–348. https://doi.org/10.1002/j.1460-2075.1994.tb06267.x

Qiao, A., Han, S., Li, X., Li, Z., Zhao, P., Dai, A., Chang, R., Tai, L., Tan, Q., Chu, X., Ma, L., Thorsen, T. S., Reedtz-Runge, S., Yang, D., Wang, M. W., Sexton, P. M., Wootten, D., Sun, F., Zhao, Q., & Wu, B. (2020). Structural basis of Gs and Gi recognition by the human glucagon receptor. Science, 367(6484), 1346–1352. https://doi.org/10.1126/science.aaz5346

Rai, D., Akagi, T., Shimohata, A., Ishii, T., Gangi, M., Maruyama, T., Wada-Kiyama, Y., Ogiwara, I., & Kaneda, M. (2021). Involvement of the C-terminal domain in cell surface localization and G-protein coupling of mGluR6. J Neurochem, 158(4), 837–848. https://doi.org/10.1111/jnc.15217

Rasmussen, S. G., Choi, H. J., Fung, J. J., Pardon, E., Casarosa, P., Chae, P. S., Devree, B. T., Rosenbaum, D. M., Thian, F. S., Kobilka, T. S., Schnapp, A., Konetzki, I., Sunahara, R. K., Gellman, S. H., Pautsch, A., Steyaert, J., Weis, W. I., & Kobilka, B. K. (2011). Structure of a nanobody-stabilized active state of the beta(2) adrenoceptor. Nature, 469(7329), 175–180. https://doi.org/10.1038/nature09648

Rasmussen, S. G., Choi, H. J., Rosenbaum, D. M., Kobilka, T. S., Thian, F. S., Edwards, P. C., Burghammer, M., Ratnala, V. R., Sanishvili, R., Fischetti, R. F., Schertler, G. F., Weis, W. I., & Kobilka, B. K. (2007). Crystal structure of the human beta2 adrenergic G-protein-coupled receptor. Nature, 450(7168), 383–387. https://doi.org/10.1038/nature06325

Rasmussen, S. G., DeVree, B. T., Zou, Y., Kruse, A. C., Chung, K. Y., Kobilka, T. S., Thian, F. S., Chae, P. S., Pardon, E., Calinski, D., Mathiesen, J. M., Shah, S. T., Lyons, J. A., Caffrey, M., Gellman, S. H., Steyaert, J., Skiniotis, G., Weis, W. I., Sunahara, R. K., & Kobilka, B. K. (2011). Crystal structure of the beta2 adrenergic receptor-Gs protein complex. Nature, 477(7366), 549–555. https://doi.org/10.1038/nature10361

Rondard, P., Goudet, C., Kniazeff, J., Pin, J. P., & Prezeau, L. (2011). The complexity of their activation mechanism opens new possibilities for the modulation of mGlu and GABAB class C G protein-coupled receptors. Neuropharmacology, 60(1), 82–92. https://doi.org/10.1016/j.neuropharm.2010.08.009

Rose, A. S., Elgeti, M., Zachariae, U., Grubmuller, H., Hofmann, K. P., Scheerer, P., & Hildebrand, P. W. (2014). Position of transmembrane helix 6 determines receptor G protein coupling specificity. J Am Chem Soc, 136(32), 11244–11247. https://doi.org/10.1021/ja5055109

Rose, A. S., Zachariae, U., Grubmuller, H., Hofmann, K. P., Scheerer, P., & Hildebrand, P. W. (2015). Role of Structural Dynamics at the Receptor G Protein Interface for Signal Transduction. PLoS One, 10(11), e0143399. https://doi.org/10.1371/journal.pone.0143399

Rosenbaum, D. M., Rasmussen, S. G., & Kobilka, B. K. (2009). The structure and function of G-protein-coupled receptors. Nature, 459(7245), 356–363. https://doi.org/10.1038/nature08144

Rosenbaum, D. M., Zhang, C., Lyons, J. A., Holl, R., Aragao, D., Arlow, D. H., Rasmussen, S. G., Choi, H. J., Devree, B. T., Sunahara, R. K., Chae, P. S., Gellman, S. H., Dror, R. O., Shaw, D. E., Weis, W. I., Caffrey, M., Gmeiner, P., & Kobilka, B. K. (2011). Structure and function of an irreversible agonist-beta(2) adrenoceptor complex. Nature, 469(7329), 236–240. https://doi.org/10.1038/nature09665

Salom, D., Lodowski, D. T., Stenkamp, R. E., Le Trong, I., Golczak, M., Jastrzebska, B., Harris, T., Ballesteros, J. A., & Palczewski, K. (2006). Crystal structure of a photoactivated deprotonated intermediate of rhodopsin. Proc Natl Acad Sci U S A, 103(44), 16123–16128. https://doi.org/10.1073/pnas.0608022103

Sandhu, M., Touma, A. M., Dysthe, M., Sadler, F., Sivaramakrishnan, S., & Vaidehi, N. (2019). Conformational plasticity of the intracellular cavity of GPCR-G-protein complexes leads to G-protein promiscuity and selectivity. Proc Natl Acad Sci U S A, 116(24), 11956–11965. https://doi.org/10.1073/pnas.1820944116

Seven, A. B., Barros-Alvarez, X., de Lapeyriere, M., Papasergi-Scott, M. M., Robertson, M. J., Zhang, C., Nwokonko, R. M., Gao, Y., Meyerowitz, J. G., Rocher, J. P., Schelshorn, D., Kobilka, B. K., Mathiesen, J. M., & Skiniotis, G. (2021). G-protein activation by a metabotropic glutamate receptor. Nature, 595(7867), 450–454. https://doi.org/10.1038/s41586-021-03680-3

Siuda, E. R., McCall, J. G., Al-Hasani, R., Shin, G., Il Park, S., Schmidt, M. J., Anderson, S. L., Planer, W. J., Rogers, J. A., & Bruchas, M. R. (2015). Optodynamic simulation of beta-adrenergic receptor signalling. Nat Commun, 6, 8480. https://doi.org/10.1038/ncomms9480

Spoida, K., Eickelbeck, D., Karapinar, R., Eckhardt, T., Mark, M. D., Jancke, D., Ehinger, B. V., Konig, P., Dalkara, D., Herlitze, S., & Masseck, O. A. (2016). Melanopsin Variants as Intrinsic Optogenetic On and Off Switches for Transient versus Sustained Activation of G Protein Pathways. Curr Biol, 26(9), 1206–1212. https://doi.org/10.1016/j.cub.2016.03.007

Standfuss, J., Edwards, P. C., D’Antona, A., Fransen, M., Xie, G., Oprian, D. D., & Schertler, G. F. (2011). The structural basis of agonist-induced activation in constitutively active rhodopsin. Nature, 471(7340), 656–660. https://doi.org/10.1038/nature09795

Terakita, A., Yamashita, T., Nimbari, N., Kojima, D., & Shichida, Y. (2002). Functional interaction between bovine rhodopsin and G protein transducin. J Biol Chem, 277(1), 40–46. https://doi.org/10.1074/jbc.M104960200

Tian, L., & Kammermeier, P. J. (2006). G protein coupling profile of mGluR6 and expression of G alpha proteins in retinal ON bipolar cells. Vis Neurosci, 23(6), 909–916. https://doi.org/10.1017/S0952523806230268

Tichy, A. M., So, W. L., Gerrard, E. J., & Janovjak, H. (2022). Structure-guided optimization of light-activated chimeric G-protein-coupled receptors. Structure. https://doi.org/10.1016/j.str.2022.04.012

Tsai, C. J., Marino, J., Adaixo, R., Pamula, F., Muehle, J., Maeda, S., Flock, T., Taylor, N. M., Mohammed, I., Matile, H., Dawson, R. J., Deupi, X., Stahlberg, H., & Schertler, G. (2019). Cryo-EM structure of the rhodopsin-Galphai-betagamma complex reveals binding of the rhodopsin C-terminal tail to the gbeta subunit. eLife, 8. https://doi.org/10.7554/eLife.46041

Tsai, C. J., Pamula, F., Nehme, R., Muhle, J., Weinert, T., Flock, T., Nogly, P., Edwards, P. C., Carpenter, B., Gruhl, T., Ma, P., Deupi, X., Standfuss, J., Tate, C. G., & Schertler, G. F. X. (2018). Crystal structure of rhodopsin in complex with a mini-Go sheds light on the principles of G protein selectivity. Sci Adv, 4(9), eaat7052. https://doi.org/10.1126/sciadv.aat7052

Valdez-Lopez, J. C., Petr, S. T., Donohue, M. P., Bailey, R. J., Gebreeziabher, M., Cameron, E. G., Wolf, J. B., Szalai, V. A., & Robinson, P. R. (2020). The C-Terminus and Third Cytoplasmic Loop Cooperatively Activate Mouse Melanopsin Phototransduction. Biophys J, 119(2), 389–401. https://doi.org/10.1016/j.bpj.2020.06.013

van Wyk, M., Pielecka-Fortuna, J., Lowel, S., & Kleinlogel, S. (2015). Restoring the ON Switch in Blind Retinas: Opto-mGluR6, a Next-Generation, Cell-Tailored Optogenetic Tool. PLoS Biol, 13(5), e1002143. https://doi.org/10.1371/journal.pbio.1002143

Venkatakrishnan, A. J., Deupi, X., Lebon, G., Heydenreich, F. M., Flock, T., Miljus, T., Balaji, S., Bouvier, M., Veprintsev, D. B., Tate, C. G., Schertler, G. F., & Babu, M. M. (2016). Diverse activation pathways in class A GPCRs converge near the G-protein-coupling region. Nature, 536(7617), 484–487. https://doi.org/10.1038/nature19107

Wang, C., Jiang, Y., Ma, J., Wu, H., Wacker, D., Katritch, V., Han, G. W., Liu, W., Huang, X. P., Vardy, E., McCorvy, J. D., Gao, X., Zhou, X. E., Melcher, K., Zhang, C., Bai, F., Yang, H., Yang, L., Jiang, H., … Xu, H. E. (2013). Structural basis for molecular recognition at serotonin receptors. Science, 340(6132), 610–614. https://doi.org/10.1126/science.1232807

Xiao, R. P. (2001). Beta-adrenergic signaling in the heart: dual coupling of the beta2-adrenergic receptor to G(s) and G(i) proteins. Sci STKE, 2001(104), re15. https://doi.org/10.1126/stke.2001.104.re15

Xu, F., Wu, H., Katritch, V., Han, G. W., Jacobson, K. A., Gao, Z. G., Cherezov, V., & Stevens, R. C. (2011). Structure of an agonist-bound human A2A adenosine receptor. Science, 332(6027), 322–327. https://doi.org/10.1126/science.1202793

Yamashita, T., Terakita, A., & Shichida, Y. (2001). The second cytoplasmic loop of metabotropic glutamate receptor functions at the third loop position of rhodopsin. Journal of Biochemistry, 130(1), 149–155. https://doi.org/DOI10.1093/oxfordjournals.jbchem.a002954

Yen, H. Y., Hoi, K. K., Liko, I., Hedger, G., Horrell, M. R., Song, W., Wu, D., Heine, P., Warne, T., Lee, Y., Carpenter, B., Pluckthun, A., Tate, C. G., Sansom, M. S. P., & Robinson, C. V. (2018). PtdIns(4,5)P2 stabilizes active states of GPCRs and enhances selectivity of G-protein coupling. Nature, 559(7714), 423–427. https://doi.org/10.1038/s41586-018-0325-6

Zhou, X. E., Melcher, K., & Xu, H. E. (2019). Structural biology of G protein-coupled receptor signaling complexes. Protein Sci, 28(3), 487–501. https://doi.org/10.1002/pro.3526

