## Supplemental Material for "Functional Optimization of Light-Activatable Opto-GPCRs: Illuminating the Importance of the Proximal C-terminus in G-protein Specificity"

```
mGluR2/1-872     1 -------MGSLLALLALLLLWGAVAEGPAKKVLTLEGDLVLGGLFPVHQKGGPAEDC  50
mGluR6/1-877     1 MARPRRAREPLLVALLPLAWLAQAGLARAAGSVRLAGGLTLGGLFPVHARGAAGRAC  57

mGluR2/1-872    51 GPVNEHRGIQRLEAMLFALDRINRDPHLLPGVRLGAHILDSCSKDTHALEQALDFVR 107
mGluR6/1-877    58 GQLKKEQGVHRLEAMLYALDRVNADPELLPGVRLGARLLDTCSRDTYALEQALSFVQ 114

mGluR2/1-872   108 ASLSRGADGSR--HICPDGSYATHGDAPTAITGVIGGSYSDVSIQVANLLRLFQIPQ 162
mGluR6/1-877   115 ALIRGRGDGDEVGVRCPGGVPPLRPAPPERVVAVVGASASSVSIMVANVLRLFAIPQ 171

mGluR2/1-872   163 ISYASTSAKLSDKSRYDYFARTVPPDFFQAKAMAEILRFFNWTYVSTVASEGDYGET 219
mGluR6/1-877   172 ISYASTAPELSDSTRYDFFSRVVPPDSYQAQAMVDIVRALGWNYVSTLASEGNYGES 228

mGluR2/1-872   220 GIEAF-ELEARARNICVATSEKVGRAMSRAAFEGVVRALLQKPSARVAVLFTRSEDA 275
mGluR6/1-877   229 GVEAFVQISREAGGVCIAQSIKIPREPKPGEFSKVIRRLMETPNARGIIIFANEDDI 285

mGluR2/1-872   276 RELLAASQR--LNASFTWVASDGWGALESVVAGSEGAAEGAITIELASYPISDFASY 330
mGluR6/1-877   286 RRVLEAARQANLTGHFLWVGSDSWGAKTSPILSLEDVAVGAITILPKRASIDGFDQY 342

mGluR2/1-872   331 FQSLDPWNNSRNPWFREFWEQRFRCSFRQ----------RDCAAHSL--RAVPFEQES 376
mGluR6/1-877   343 FMTRSLENNRRNIWFAEFWEENFNCKLTSSGTQSDDSTRKCTGEERIGRDSTYEQEG 399

mGluR2/1-872   377 KIMFVVNAVYAMAHALHNMHRALCPNTTRLCDAMRPVNGRRLYKDFVLNVKFDAPFR 433
mGluR6/1-877   400 KVQFVIDAVYAIAHALHSMHQALCPGHTGLCPAMEPTDGRMLLQ-YIRAVRFNGS-- 453

mGluR2/1-872   434 PADTHNEVRFDRFGDGIGRYNIFTYLRAGSGRYR--YQKVGYWAEGLTLDTSLIPWA 488
mGluR6/1-877   454 ---AGTPVMFNENGDAPGRYDIFQYQATNGSASSGGYQAVGQWAETLRLDVEALQWS 507

mGluR2/1-872   489 SPSAGPLPASRCSEPCLQNEVKSVQPGEVCCWLCIPCQPYEYRLDEFTCADCGLGYW 545
mGluR6/1-877   508 G-DPHEVPSSLCSLPCGPGERKKMVKGVPCCWHCEACDGYRFQVDEFTCEACPGDMR 563

mGluR2/1-872   546 PNASLTGCFELPQEYIRWGDAWAVGPVTIACLGALATLFVLGVFVRHNATPVVKASG 602
mGluR6/1-877   564 PTPNHTGCRPTPVVRLSWSSPWAAPPLLLAVLGIVATTTVVATFVRYNNTPIVRASG 620

mGluR2/1-872   603 RELCYILLGGVFLCYCMTFIFIAKPSTAVCTLRRLGLGTAFSVCYSALLTKTNRIAR 659
mGluR6/1-877   621 RELSYVLLTGIFLIYAITFLMVAEPGAAVCAARRLFLGLGTTLSYSALLTKTNRIYR 677

mGluR2/1-872   660 IFGGAREGAQRPRFISPASQVAICLALISGQLLIVVAWLVVEAPGT------GKETA 710
mGluR6/1-877   678 IFEQGKRSVTPPPFISPTSQLVITFSLTSLQVVGMIAWLGARPPHSVIDYEEQRTVD 734

mGluR2/1-872   711 PERREVVTLRCNHRDASMLGSLAYNVLLIALCTLYAFKTRKCPENFNEAKFIGFTMY 767
mGluR6/1-877   735 PEQARGV-LKCDMSDLSLIGCLGYSLLLMVTCTVYAIKARGVPETFNEAKPIGFTMY 790

mGluR2/1-872   768 TTCIIWLAFLPIFYVTSSDYR---VQTTTMCVSVSLSGSVVLGCLFAPKLHIILFQP 821
mGluR6/1-877   791 TTCIIWLAFVPIFFGTAQSAEKIYIQTTTLTVSLSLSASVSLGMLYVPKTYVILFHP 847

mGluR2/1-872   822 QKNVVSHRAPTSRFGSAAARASSSLGQGSGSQFVPTVCNGREVVDSTTSSL       872
mGluR6/1-877   848 EQNVQKRKRS---LKATSTVAAPPKGEDAEAHK------------------       877
```

**Suppl. Fig. S1: Sequence alignment of mGluR2 and mGluR6.** Arrows indicate the cutting sites for Mela-mGluR6 chimera design, the hypervariable IL3 region (between the two red arrows), the proximal C-terminus (longCT, blue arrow) and the distal C-terminus (shortCT, green arrow).

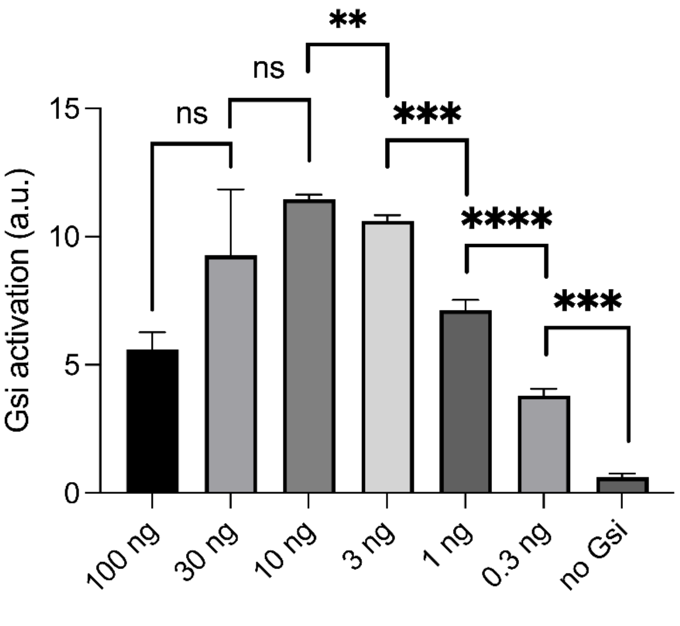


**Suppl. Fig. S2: GsX assay calibration: optimization of the amount of G-protein plasmid to be transfected into HEK293 cells.** HEK293 WT cells transfected with rhodopsin plasmid, GloSensor reporter and the different amounts of C-terminal mutated Gsi G-protein DNA to determine the optimal rhodopsin-to-Gsi-protein ratio for maximal luminescence. All DNA concentrations were tested in the same plate, baseline luminescence subtracted and the achieved max. cAMP value for each concentration of Gsi-protein is shown here. We found the 10:1 ratio of plasmid:GsX to be optimal. N=3

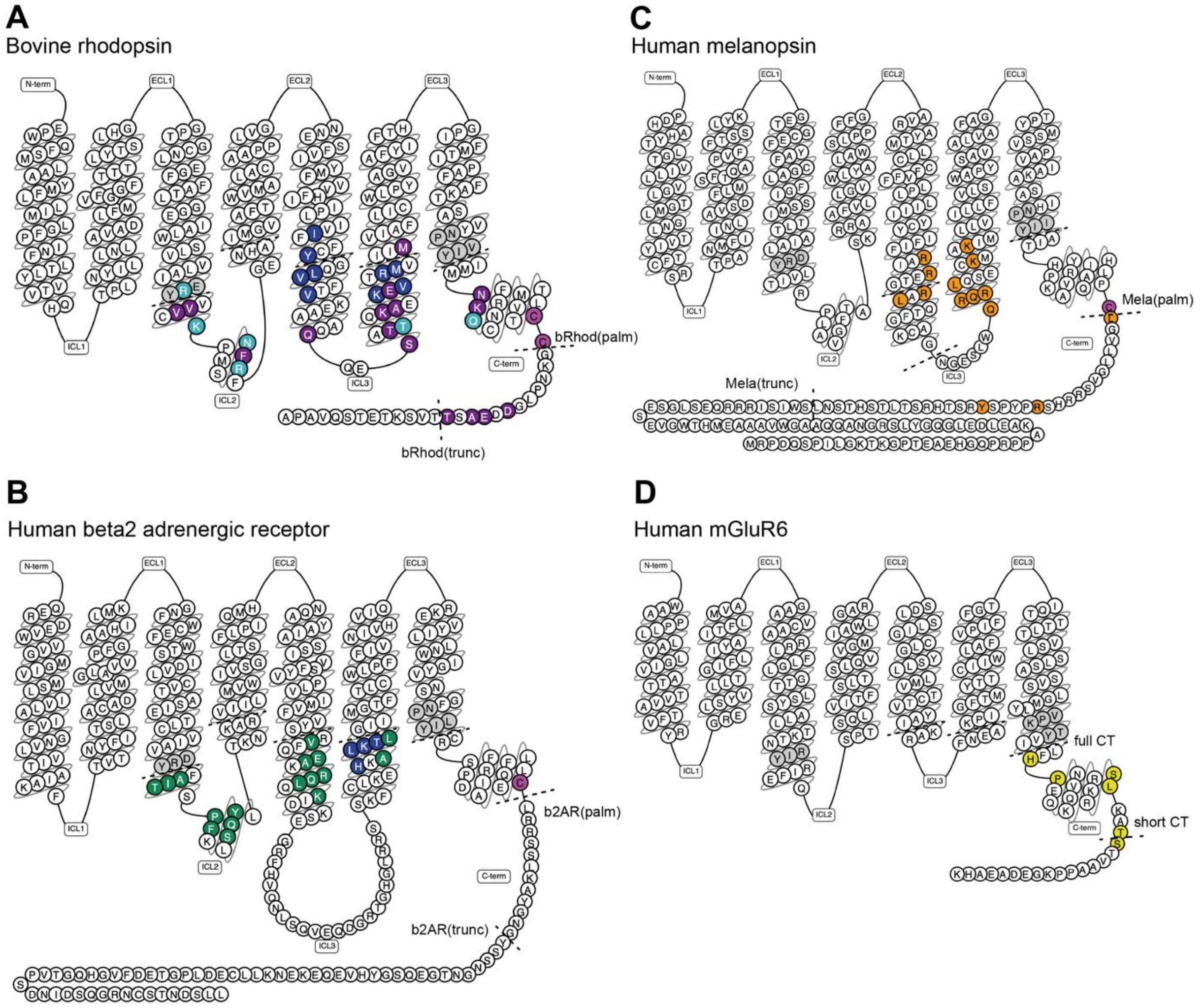


**Suppl. Fig. S3: Snake plots of the four GPCRs employed, highlighting known G-protein contact residues as well as the used chimeric recombination sites.** (A) bovine rhodopsin (B) human beta-2 adrenergic receptor (C) human melanopsin (D) human mGluR6. Highlighted are residues that have been shown to be in proximity to or form residue-residue contacts with the G-protein: purple (Tsai et al., 2019), light blue (Kang et al., 2018), dark blue (Zhou et al., 2019), green (Rasmussen et al., 2011), orange (Valdez-Lopez et al., 2020). Highlighted in yellow are residues involved in cell surface localization and G-protein coupling of mGluR6 (Rai et al., 2021). Notice that albeit drawn, mGluR6 does probably not possess an ordered H8 nor a known palmitoylation site (highlighted in pink in the other plots A-C). Highlighted in grey are the conserved NPxxY (VPKTY in mGluR6) and DR(I)Y/ERY motifs at the end of TM7 and at the start of IL2, respectively. See also sequence information in Fig. 1B,D.

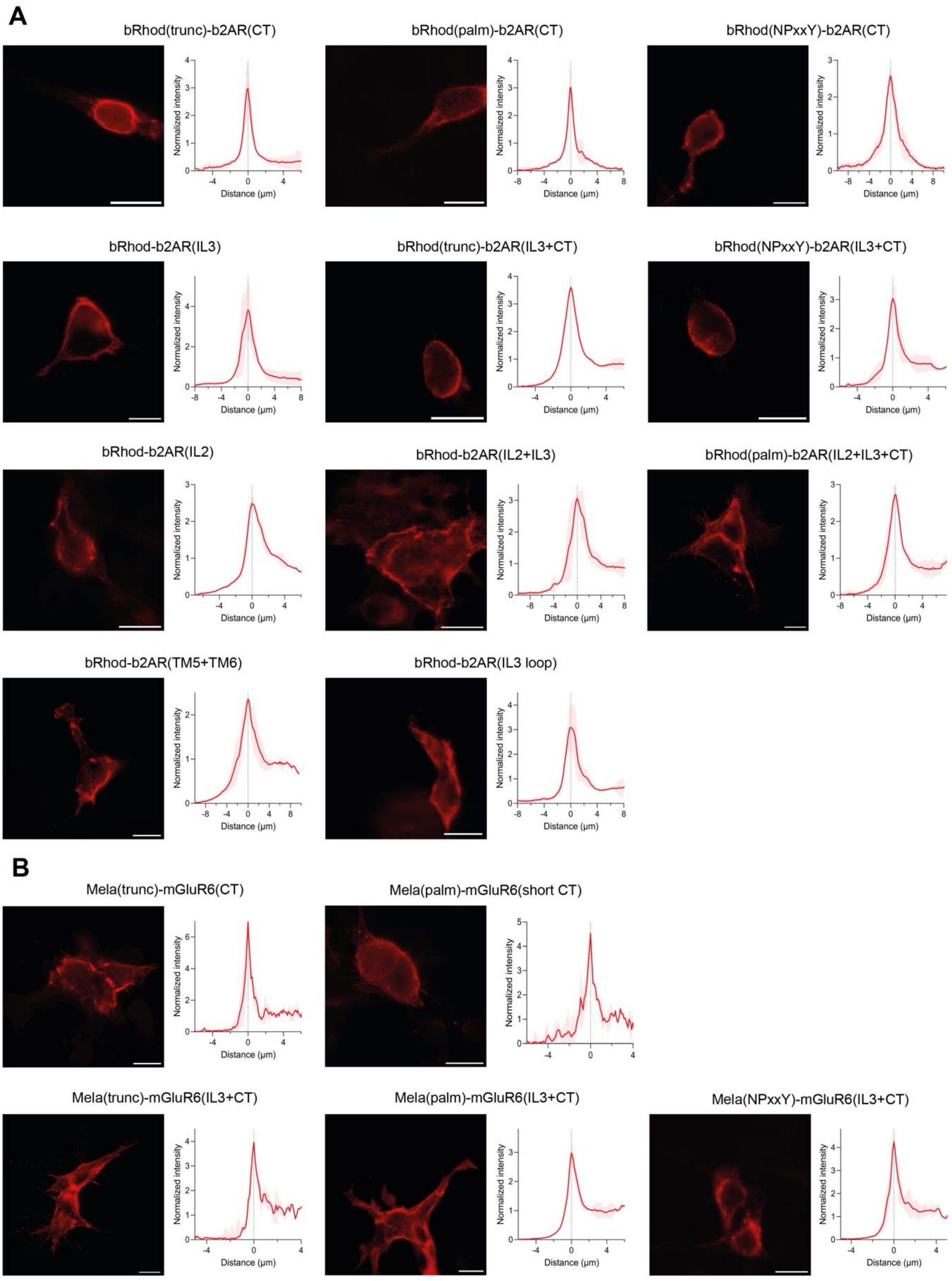


**Suppl. Fig. S4: Cell surface expression of bRhod-b2AR and melanopsin-mGluR6 chimeras.** Fluorescence images of the cell surface expression of the library of chimeras tested between bovine rhodopsin and b2AR (A) and between melanopsin and mGluR6 (B) with their respective cross-sectional normalized fluorescence intensities across the cell membrane. The dashed line at X=0 in the fluorescence intensity plots corresponds to the cell membrane across which the intensity profile was plotted. N=3. Scale bar=10µm.

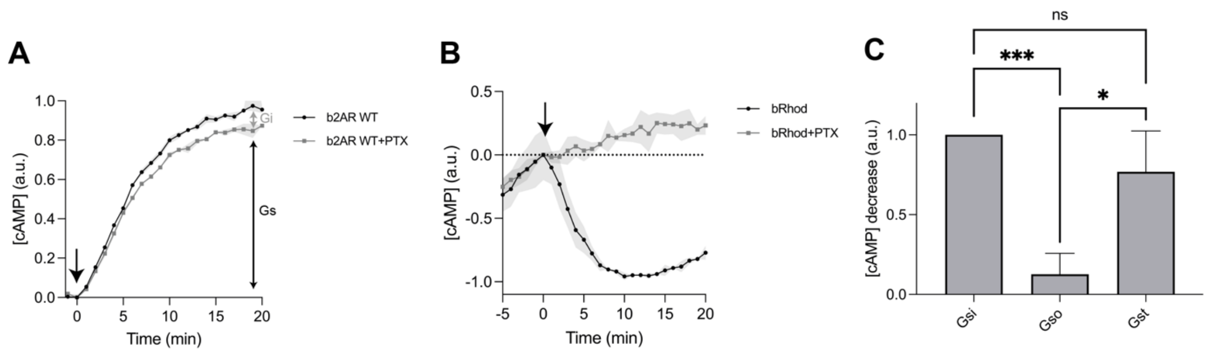


**Suppl. Fig. S5: Signaling profiles of wildtype bovine rhodopsin and beta-2 adrenergic receptor in the GloSensor second messenger assay.** (A) cAMP assay of b2AR without (black) and with addition of PTX (grey) that blocks activation of endogenous Gi proteins of HEK293 cells. This shows that b2AR almost exclusively activates Gs-proteins (92.9 ± 0.7%, black double arrow), elevating cAMP and to a much lesser degree Gi proteins (7.1 ± 0.8%, grey double arrow), lowering cAMP levels in the cell. The reporter luminescence of b2AR was normalized between 0 and 1 and the luminescence value with PTX was normalized to b2AR. (B) cAMP assay of bovine rhodopsin without (black) and with addition of PTX (grey). Intracellular cAMP levels were elevated with forskolin before application of the light stimulus at time point zero. The traces show that bovine rhodopsin is exclusively activates Gi proteins, lowering cAMP levels in the cell. The reporter luminescence levels of rhodopsin were normalized between 0 and 1 and the luminescence values with PTX were normalized to rhodopsin. Baseline was subtracted for all luminescence and values adjusted to y=0 for x=0. (C) HEK293 cells transfected with OptoGPCR plasmids, GloSensor reporter and the C-terminal mutated Gsi, Gso, or Gst G-proteins and subjected to a Gs cAMP assay. Bovine rhodopsin preferentially binds to Gst (transducing), its natural effector in retinal photoreceptors, but equally well to Gsi and to a lesser degree (12.7 ± 0.13%) to Gso. This may explain why rhodopsin functioned as an optogenetic tool in retinal-ON bipolar cells (Cehajic-Kapetanovic et al., 2015)where the cellular response is mediated by mGluR6 shown to be mainly Go selective, but also minimally Gi selective (Tian & Kammermeier, 2006). N=3; light stimulus: array of 3x8 LEDs, 470 nm, $5x10^{16}$ photons/cm2/s.

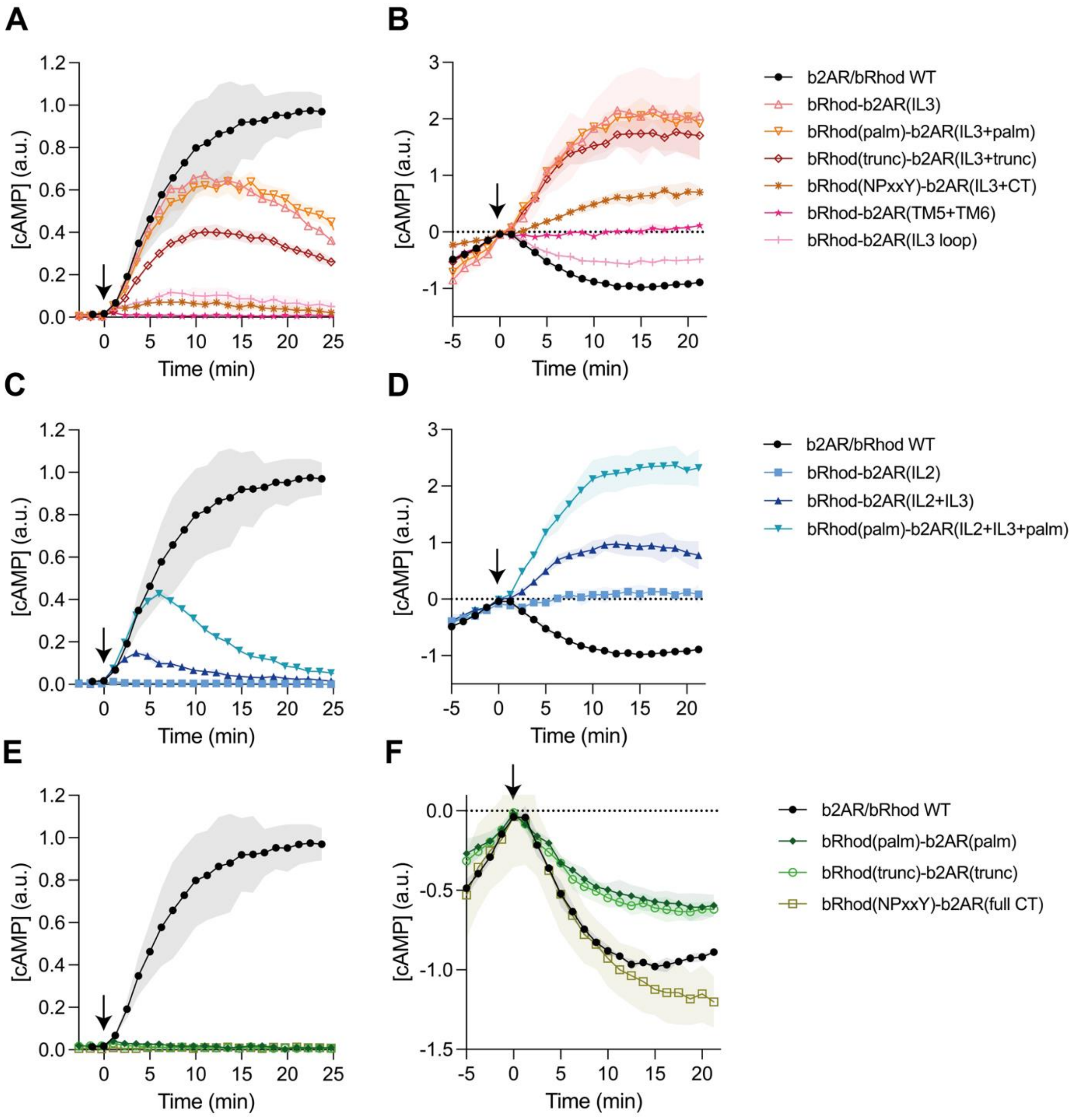


**Suppl. Fig. S6: Specificity of G-protein pathway activation of bRhod-b2AR chimeras determined with the GloSensor second messenger assay.** (A,C,E) cAMP assay for Gs activation and (B,D,F) Gi activation. Intracellular cAMP levels were elevated with forskolin before application of the light stimulus at time point zero in (B,D,F). cAMP levels were measured every 15s. The reporter luminescence levels of WT b2AR (A,C,E) or WT bRhod (B,D,F) were normalized between 0 and 1 and the luminescence values of the variants were normalized to b2AR (A,C,E) or bRhod (B,D,F). The baseline was adjusted for x=0 to equal y=0. No Gq activation (increase in Ca2+) was measured for any variant (not shown here). The timepoint of light stimulation is shown with an arrow. Please refer to the Results section and Fig. 2 of the main text for analysis and conclusions. Besides the described impact of the IL3 on Gs-signaling, shown here is the inability of the TM5+TM6 domains as well as the IL3 loop on their own to shift tropism towards Gs (A,B). Additionally, (C) and (D) show the low signaling efficacy of the IL2 replacements. N=3; light stimulus: array of 3x8 LEDs, 470 nm, $5x10^{16}$ photons/cm2/s.

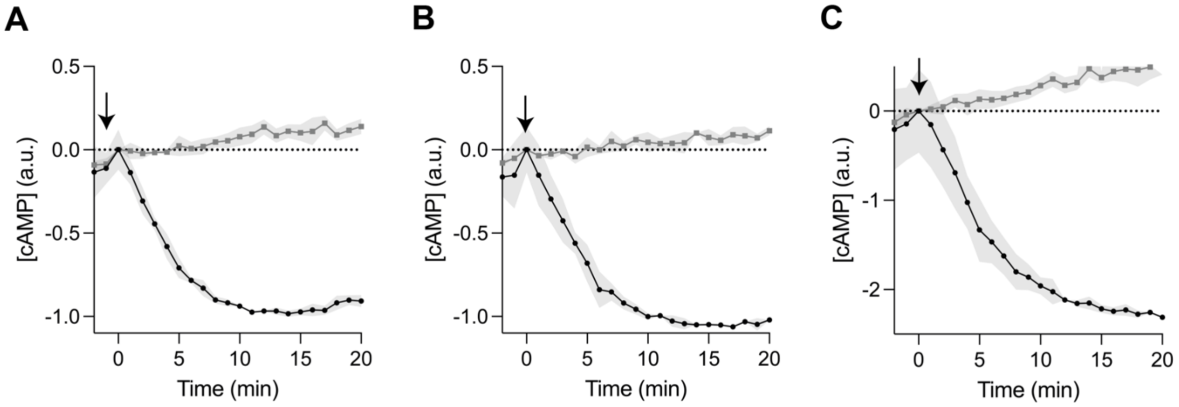


**Suppl. Fig. S7: Gi specificity of bRhod-b2AR C-terminus measured with GloSensor the second messenger assays.** (A-C) cAMP assays for Gi activation of the bRhod-b2AR C-terminal variants (A: bRhod-b2AR(trunc), B: bRhod-b2AR(palm), C: bRhod-b2AR(fullCT)) incubated overnight without (black) and with PTX (grey) that blocks endogenous Gi repsonses of HEK293 cells to uncover the pure Gs response, if existent. As obvious from the traces, all chimeras show a pure Gi response. Consequently, the C-terminus of b2AR is unable to adapt the tropism of bovine rhodopsin to the Gs protein. Intracellular cAMP levels were elevated with forskolin before application of the light stimulus at time point zero (arrow). The GloSensor reporter luminescence values of WT bRhod were normalized between 0 and 1 and the luminescence values of the variants were normalized to WT bRhod. The baseline was adjusted for x=0 to equal y=0. N=3; light stimulus: array of 3x8 LEDs, 470 nm, $5x10^{16}$ photons/$cm^2$/s.

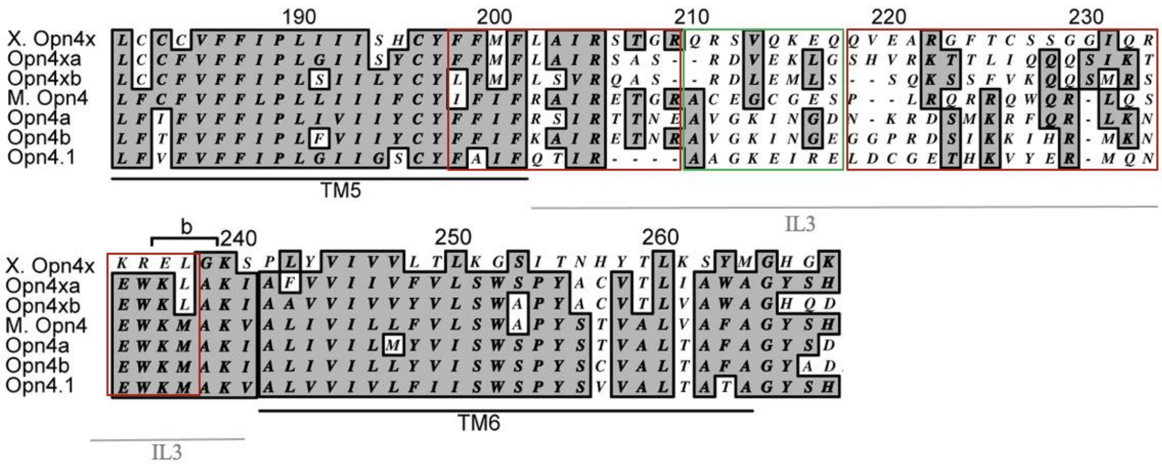


**Suppl. Fig. S8: Alignment of melanopsin genes of different species.** Alignment of melanopsin genes (OPN4) from different species indicating the hypervariable regions in IL3 between TM5 and TM6 (Matos-Cruz et al., 2011). Indicated in green is the cytoplasmic loop region, which we replaced with the analogous region of mGluR6. It is obvious that the sequence variability also extends into the TM5 and TM6 cytoplasmic protrusions (indicated in red).

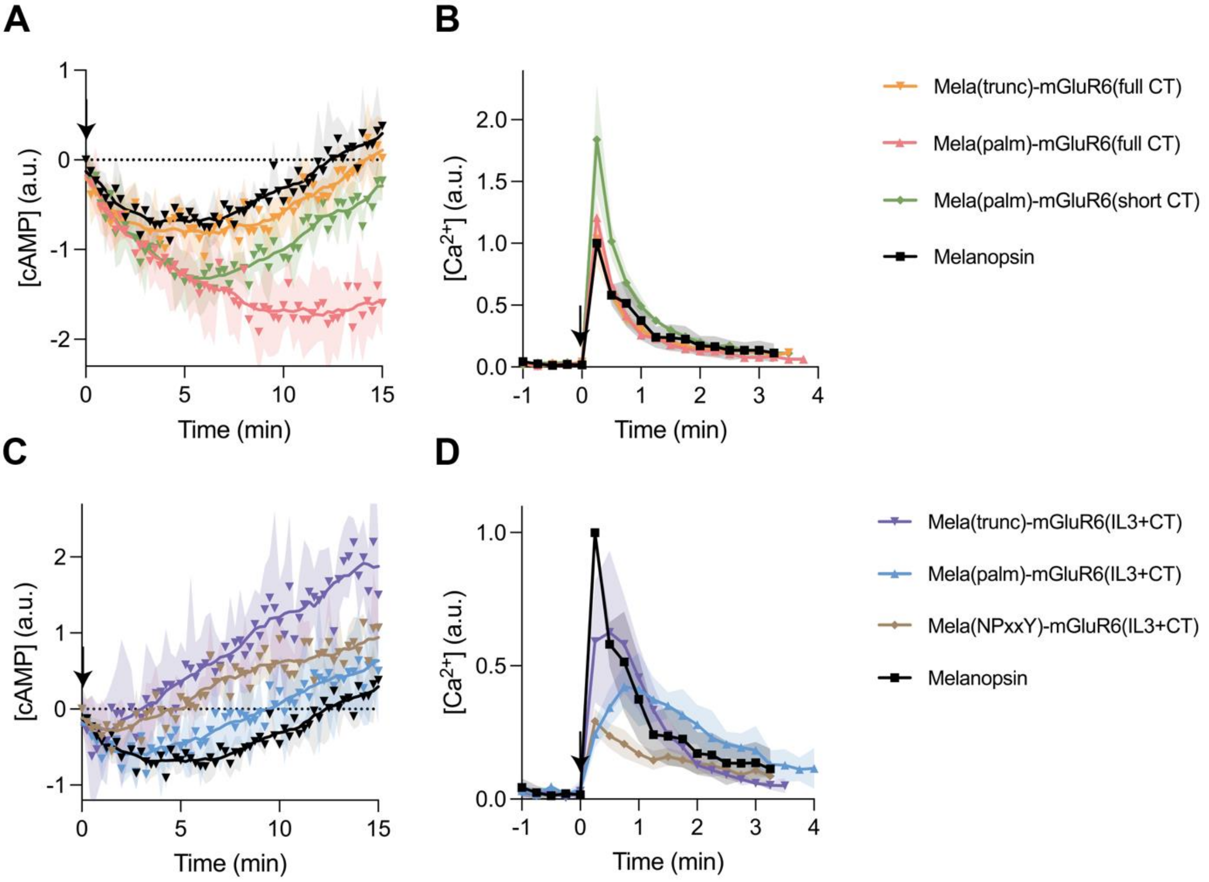


**Suppl. Fig. S9: Specificity of G-protein pathway activation of Mela-mGluR6 chimeras determined with the GloSensor second messenger assay.** (A,C) cAMP assay for Gi activation and (B,D) Ca2+ assay for Gq activation of the (A,B) Mela-mGluR6 C-terminal variants and the (C,D) Mela-mGluR6 IL3+CT variants. Intracellular cAMP levels were elevated with forskolin before application of the light stimulus at time point zero in (A,C). The reporter luminescence levels of WT melanopsin were normalized between 0 and 1 and the luminescence values of the variants were normalized to melanopsin. The baseline was adjusted for x=0 to equal y=0. No Gs activation (increase in cAMP) was measured for any variant (not shown here). Compare this raw data with the analysed data in Figs 3 & 4 in the main manuscript. N=9; statistical significance: * = $p<0.05$, ** = $p<0.01$; light stimulus: array of 3x8 LEDs, 470 nm, $5x10^{16}$ photons/cm$^2$/s.

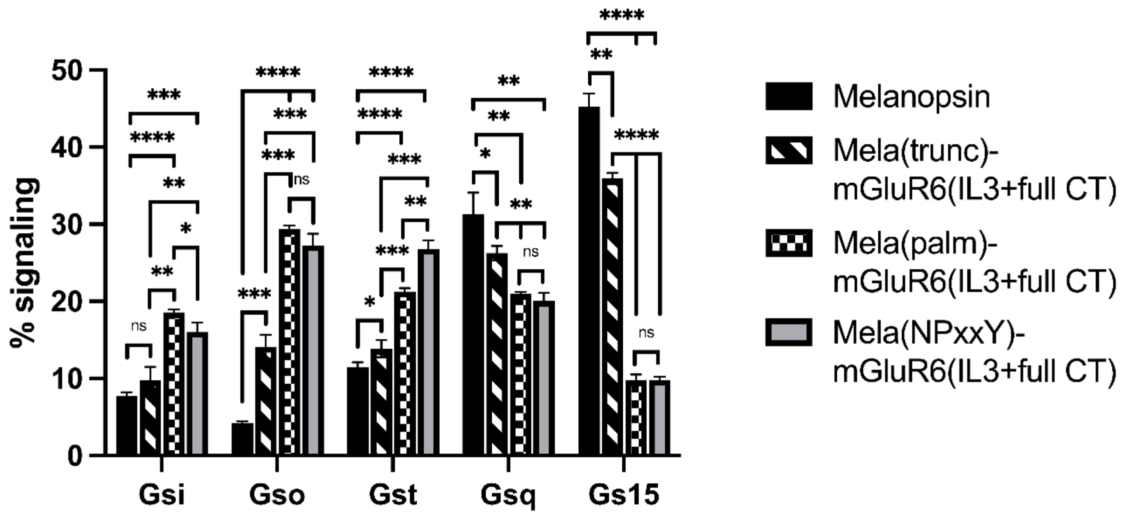


**Suppl. Fig. S10: Specificity of G-protein pathway activation of Mela-mGluR6 IL3+CT chimeras determined with the GsX GloSensor second messenger assays.** HEK293 WT cells transfected with the respective OptoGPCR plasmid, GloSensor reporter and the C-terminal mutated Gsq, Gs15, Gsi, Gso, or Gst G-proteins and subjected to a Gs cAMP assay. All GloSensor cAMP luminescence values were normalized to mKate2 fluorescence (to normalize to the number of cells measured) and the baseline fluorescence was subtracted. The percentual proportion of each G-protein subfamily in total Gq and Gi signaling (100%) was calculated by dividing the max. cAMP response of each GsX-protein by the total sum of all max. GsX values. The influence mediated by the proximal C-terminus (H8 palmitoylation site at C367 to AA400 in shifting tropism from mainly Gq to mainly Gi is evident in this Gi and Gq subfamily signaling fingerprint. This can be seen in 1) the significant increase in Gsi, Gso and Gst signaling between the Mela(trunc)-mGluR6(IL3+CT) and the Mela(palm)-mGluR6(IL3+CT) and 2) the significant decrease in Gsq and Gs15 signaling between these variants. See also Fig. 3 and main text. N=3; light stimulus: array of 3x8 LEDs, 470 nm, $5x10^{16}$ photons/$cm^2$/s.

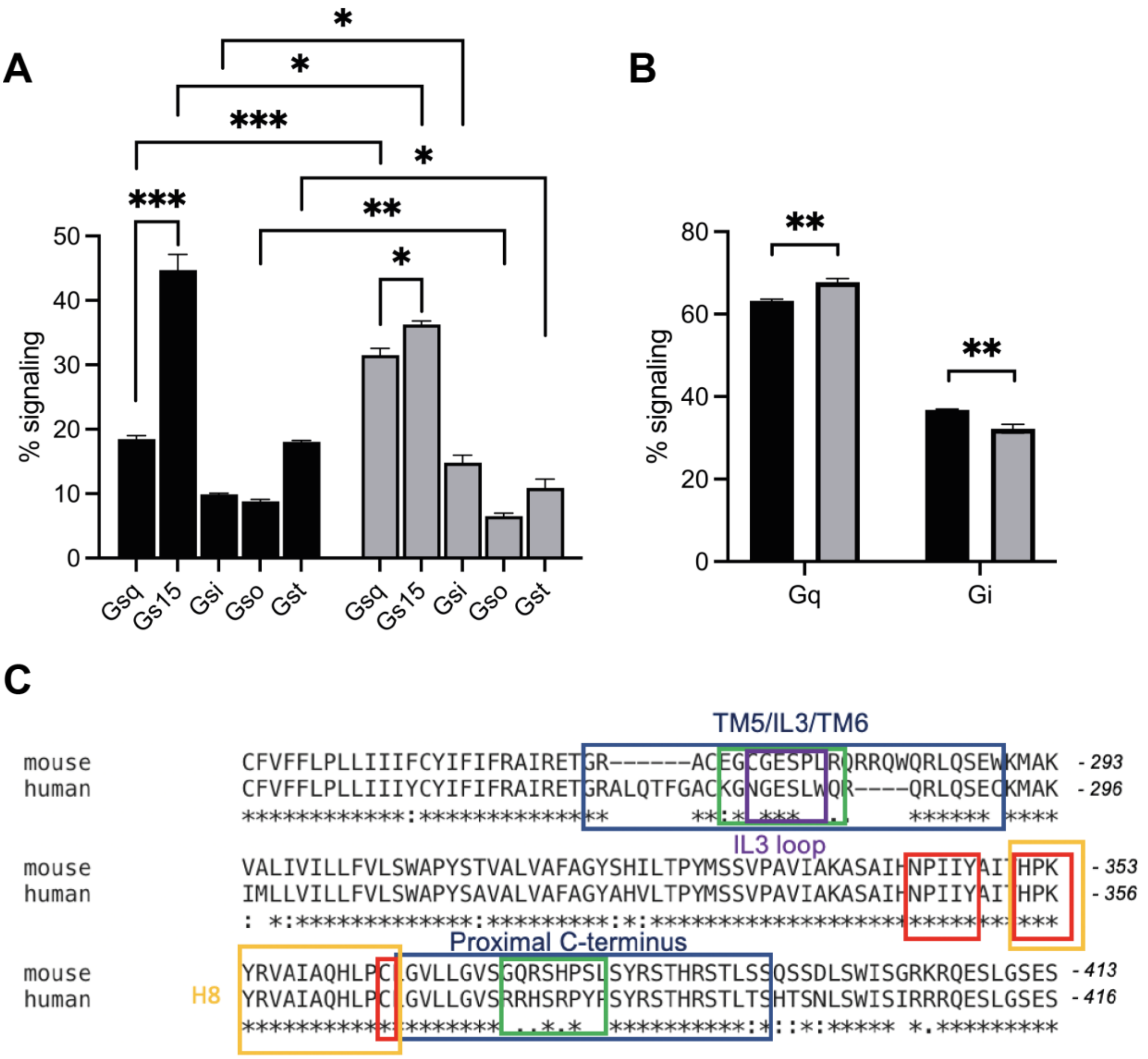


**Suppl. Fig. S11: Gi and Gq signaling fingerprints of human and murine melanopsin highlighting the differences.** HEK293 WT cells transfected with human (black) or mouse (grey) melanopsin. All GloSensor cAMP luminescence values were normalized to mKate2 fluorescence (e.g. number of cells measured) and the baseline subtracted. (A) The percentual proportion of each G-protein subfamily in total Gq and Gi signaling (100%) was calculated by dividing the max. cAMP response of each GsX-protein by the total sum of all max. GsX values. Albeit melanopsin from both species favourably signal via G15, murine melanopsin shows a much more balanced Gq and G15 response. Human melanopsin, however, shows a preference of Gi and Gt signaling compared to murine melanopsin. (B) The total Gq and Gi signaling was calculated by adding the max. cAMP value of the sum of Gsq and Gs15, and Gsi, Gso, and Gst, respectively, and dividing by the sum of all GsX responses to determine the percentage of the two main G-protein families in total signaling. The panel shows that while murine melanopsin signals significantly stronger via Gq, whereas human melanopsin signals significantly stronger via Gi compared to murine melanopsin. N=3; light stimulus: 470 nm, $5x10^{16}$ photons/cm$^2$/s. (C) Sequence comparison of human and murine melanopsin. Interestingly, the two sequences diverge exactly in the IL3 and the proximal C-terminal regions (boxed in green) investigate in this study and postulated to be involved in G-protein selectivity. (TM5/IL3/TM6 and the proximal C-terminus boxed in dark blue). Conservation levels are shown as *, : and . . Conserved motifs, such as the NPxxY motif at the end of TM8, the HPK motif at the start of H8 and the palmitoylated C367 at the end of H8 are highlighted in red. H8 is indicated in yellow and the cytoplasmic IL3 loop region in purple.

**Table 1: Overview of chimeric constructs**

| Construct name | Cutting site(s) opsin | Cutting site(s) target receptor |
|---|---|---|
| Melanopsin-mGluR6 | Melanopsin | mGluR6 |
| Mela(trunc)-mGluR6(CT) | Distal CT truncation site | Full CT |
| **Mela(palm)-mGluR6(CT)** | **CT palmitoylation site** | **Full CT** |
| Mela(palm)-mGluR6(short CT) | CT palmitoylation site | After distal CT truncation site |
| Mela(trunc)-mGluR6(IL3+CT) | IL3: intracellular hypervariable loop region<br>CT: distal truncation site | IL3: intracellular hypervariable loop region<br>CT: full CT |
| Mela(palm)-mGluR6(IL3+CT) | IL3: intracellular hypervariable loop region<br>CT: palmitoylation site | IL3: intracellular hypervariable loop region<br>CT: full CT |
| Mela(NPxxY)-mGlur6(IL3+CT) | IL3: intracellular hypervariable loop region<br>CT: after NPxxY | IL3: intracellular hypervariable loop region<br>CT: full CT |
| bRhod-b2AR | Bovine rhodopsin | Beta2-adrenergic receptor |
| bRhod(trunc)-b2AR(CT) | CT distal truncation site | CT distal truncation site |
| bRhod(palm)-b2AR(CT) | CT palmitoylation site | CT palmitoylation site |
| bRhod(NPxxY)b2AR(CT) | After NPxxY | After NPxxY |
| bRhod-b2AR(IL3) | TM5-IL3-TM6 | TM5-IL3-TM6 |
| bRhod-b2AR(IL3 loop) | Intracellular IL3 loop | Intracellular IL3 loop |
| bRhod(trunc)-b2AR(IL3+CT) | IL3: TM5-IL3-TM6<br>CT: distal truncation site | IL3: TM5-IL3-TM6<br>CT: distal truncation site |
| **bRhod(palm)-b2AR(IL3+CT)** | **IL3: TM5-IL3-TM6<br>CT: palmitoylation site** | **IL3: TM5-IL3-TM6<br>CT: palmitoylation site** |
| bRhod(NPxxY)-b2AR(IL3+CT) | IL3: TM5-IL3-TM6<br>CT: after NPxxY | IL3: TM5-IL3-TM6<br>CT: after NPxxY |
| bRhod-b2AR(IL2) | TM3-IL2-TM4 | TM3-IL2-TM4 |
| bRhod-b2AR(IL2+IL3) | IL2: TM3-IL2-TM4<br>IL3: TM5-IL3-TM6 | IL2: TM3-IL2-TM4<br>IL3: TM5-IL3-TM6 |
| bRhod-b2AR(IL2+IL3+CT) | IL2: TM3-IL2-TM4<br>IL3: TM5-IL3-TM6<br>CT: palmitoylation site | IL2: TM3-IL2-TM4<br>IL3: TM5-IL3-TM6<br>CT: palmityolation site |
| bRhod-b2AR(TM5+TM6) | TM5+TM6 without IL3 loop | TM5+TM6 without IL3 loop |

Overview of the chimeras with naming and the respective cutting sites in the opsin and target receptor pairs (melanopsin and mGluR6, and bovine rhodopsin and b2AR). The functionally optimized and most promising constructs are indicated in bold, highlighting the necessary replacements to induce the desired downstream signaling pathway of the target receptor. Please refer to Fig. 1A for cutting sites.

**Literature**


Cehajic-Kapetanovic, J., Eleftheriou, C., Allen, A. E., Milosavljevic, N., Pienaar, A., Bedford, R., Davis, K. E., Bishop, P. N., & Lucas, R. J. (2015). Restoration of Vision with Ectopic Expression of Human Rod Opsin. *Curr Biol*, *25*(16), 2111-2122. https://doi.org/10.1016/j.cub.2015.07.029

Kang, Y., Kuybeda, O., de Waal, P. W., Mukherjee, S., Van Eps, N., Dutka, P., Zhou, X. E., Bartesaghi, A., Erramilli, S., Morizumi, T., Gu, X., Yin, Y., Liu, P., Jiang, Y., Meng, X., Zhao, G., Melcher, K., Ernst, O. P., Kossiakoff, A. A., . . . Xu, H. E. (2018). Cryo-EM structure of human rhodopsin bound to an inhibitory G protein. *Nature*, *558*(7711), 553-558. https://doi.org/10.1038/s41586-018-0215-y

Matos-Cruz, V., Blasic, J., Nickle, B., Robinson, P. R., Hattar, S., & Halpern, M. E. (2011). Unexpected diversity and photoperiod dependence of the zebrafish melanopsin system. *PLoS One*, *6*(9), e25111. https://doi.org/10.1371/journal.pone.0025111

Rai, D., Akagi, T., Shimohata, A., Ishii, T., Gangi, M., Maruyama, T., Wada-Kiyama, Y., Ogiwara, I., & Kaneda, M. (2021). Involvement of the C-terminal domain in cell surface localization and G-protein coupling of mGluR6. *J Neurochem*, *158*(4), 837-848. https://doi.org/10.1111/jnc.15217

Rasmussen, S. G., DeVree, B. T., Zou, Y., Kruse, A. C., Chung, K. Y., Kobilka, T. S., Thian, F. S., Chae, P. S., Pardon, E., Calinski, D., Mathiesen, J. M., Shah, S. T., Lyons, J. A., Caffrey, M., Gellman, S. H., Steyaert, J., Skiniotis, G., Weis, W. I., Sunahara, R. K., & Kobilka, B. K. (2011). Crystal structure of the beta2 adrenergic receptor-Gs protein complex. *Nature*, *477*(7366), 549-555. https://doi.org/10.1038/nature10361

Tian, L., & Kammermeier, P. J. (2006). G protein coupling profile of mGluR6 and expression of G alpha proteins in retinal ON bipolar cells. *Vis Neurosci*, *23*(6), 909-916. https://doi.org/10.1017/S0952523806230268

Tsai, C. J., Marino, J., Adaixo, R., Pamula, F., Muehle, J., Maeda, S., Flock, T., Taylor, N. M., Mohammed, I., Matile, H., Dawson, R. J., Deupi, X., Stahlberg, H., & Schertler, G. (2019). Cryo-EM structure of the rhodopsin-Galphai-betagamma complex reveals binding of the rhodopsin C-terminal tail to the gbeta subunit. *eLife*, *8*. https://doi.org/10.7554/eLife.46041

Valdez-Lopez, J. C., Petr, S. T., Donohue, M. P., Bailey, R. J., Gebreeziabher, M., Cameron, E. G., Wolf, J. B., Szalai, V. A., & Robinson, P. R. (2020). The C-Terminus and Third Cytoplasmic Loop Cooperatively Activate Mouse Melanopsin Phototransduction. *Biophys J*, *119*(2), 389-401. https://doi.org/10.1016/j.bpj.2020.06.013

Zhou, X. E., Melcher, K., & Xu, H. E. (2019). Structural biology of G protein-coupled receptor signaling complexes. *Protein Sci*, *28*(3), 487-501. https://doi.org/10.1002/pro.3526
